# Microbiota-induced active translocation of peptidoglycan across the intestinal barrier dictates its within-host dissemination

**DOI:** 10.1101/2022.04.20.488924

**Authors:** Richard Wheeler, Paulo A. D. Bastos, Olivier Disson, Aline Rifflet, Julia Spielbauer, Marc Lecuit, Ivo Gomperts Boneca

**Author notes:** Contributed equally (Richard Wheeler, Paulo A. D. Bastos). Contributed equally (Olivier Disson, Aline Rifflet).

## Abstract

Peptidoglycan, the major structural polymer forming the cell wall of bacteria, is an important mediator of physiological and behavioral effects in mammalian hosts. These effects are frequently linked to its translocation from the intestinal lumen to host tissues. However, the modality and regulation of this translocation across the gut barrier has not been precisely addressed. In this study, we characterized the absorption of peptidoglycan across the intestine and its systemic dissemination. We report that peptidoglycan has a distinct tropism for host organs when absorbed via the gut, most notably by favoring access to the brain. We demonstrate that intestinal translocation of peptidoglycan occurs through a microbiota-induced active process. This process is regulated by the parasympathetic pathway via the muscarinic acetylcholine receptors. Together, this study reveals fundamental parameters concerning the uptake of a major microbiota molecular signal from the steady-state gut.

## Main Text

Within the intestinal tract, the gut microbiota generates signals that are recognized by different classes of host immune and non-immune cells^1^. These signals can come in the form of Microbe Associated Molecular Patterns (MAMPs), a highly diverse group of microorganism-specific molecules that are recognized by pattern recognition receptors (PRRs) expressed mainly by innate immune cells^2^. One such class of MAMP, the bacterial cell wall component peptidoglycan, is well-characterized as a key inflammatory molecule in acute and chronic conditions, and increasingly recognized as an important mediator of steady-state phenomena in the host. These effects are not limited to the gut where the microbiota resides^3^, but include the broader host physiology and behaviors, facilitated by the continuous dissemination of peptidoglycan fragments across the intestinal epithelial barrier^4–10^, for which a variety of mechanisms have been proposed^11^. Nevertheless, a clear understanding of the basic principles governing the absorption and dissemination of microbiota peptidoglycan throughout the host system is lacking.

In this study, we characterize the absorption of peptidoglycan across the gut epithelial barrier and its systemic dissemination in the steady-state. In a mouse model, we show that translocation across the intestinal barrier favors the accumulation of peptidoglycan in specific host organs, most notably the brain. We further demonstrate that the steady-state absorption of peptidoglycan is dependent on the microbial colonization status of the host. Focusing on the small intestinal epithelial barrier, we observed peptidoglycan absorption by enteric epithelial cells, among which goblet cells displayed a prominent affinity, and found that the trafficking of peptidoglycan from the gut is regulated via muscarinic acetylcholine receptors. Together, our data provide new insights into gut-axis governing the systemic presence of peptidoglycan.

## Results

We first aimed to establish the dynamics of peptidoglycan dissemination from the gut to the host system, by tracking the fate of an orally delivered, single dose of peptidoglycan in conventional adult mice. *Escherichia coli* peptidoglycan was radiolabeled by incorporation of ^3^H-*meso*DAP, and solubilized enzymatically by glycoside hydrolysis of the glycan chain to generate soluble peptidoglycan moieties (termed muropeptides; **Fig. S1A and B**). *E. coli* peptidoglycan is a relevant proxy for microbiota peptidoglycan, since the most abundant muropeptides are well represented among the structures naturally present in the intestinal microbiota peptidoglycome **(Fig. S1C)**. ^3^H-*meso*DAP labelled peptidoglycan ([^3^H]-PGN, 400,000 cpm) was administered *per os* to specific-pathogen-free (SPF) C57BL/6j mice. The blood and PBS-perfused organs were collected at time-points between 2h and 8h, processed for scintillation counting, and the [^3^H]-PGN content assessed **(Fig. 1A)**. We selected this strategy because 1) labelling with ^3^H-*meso*DAP does not alter the native composition of peptidoglycan, allowing us to track peptidoglycan undergoing natural processing within the host, and 2) scintillation counting is highly sensitive.

**Fig. 1.**
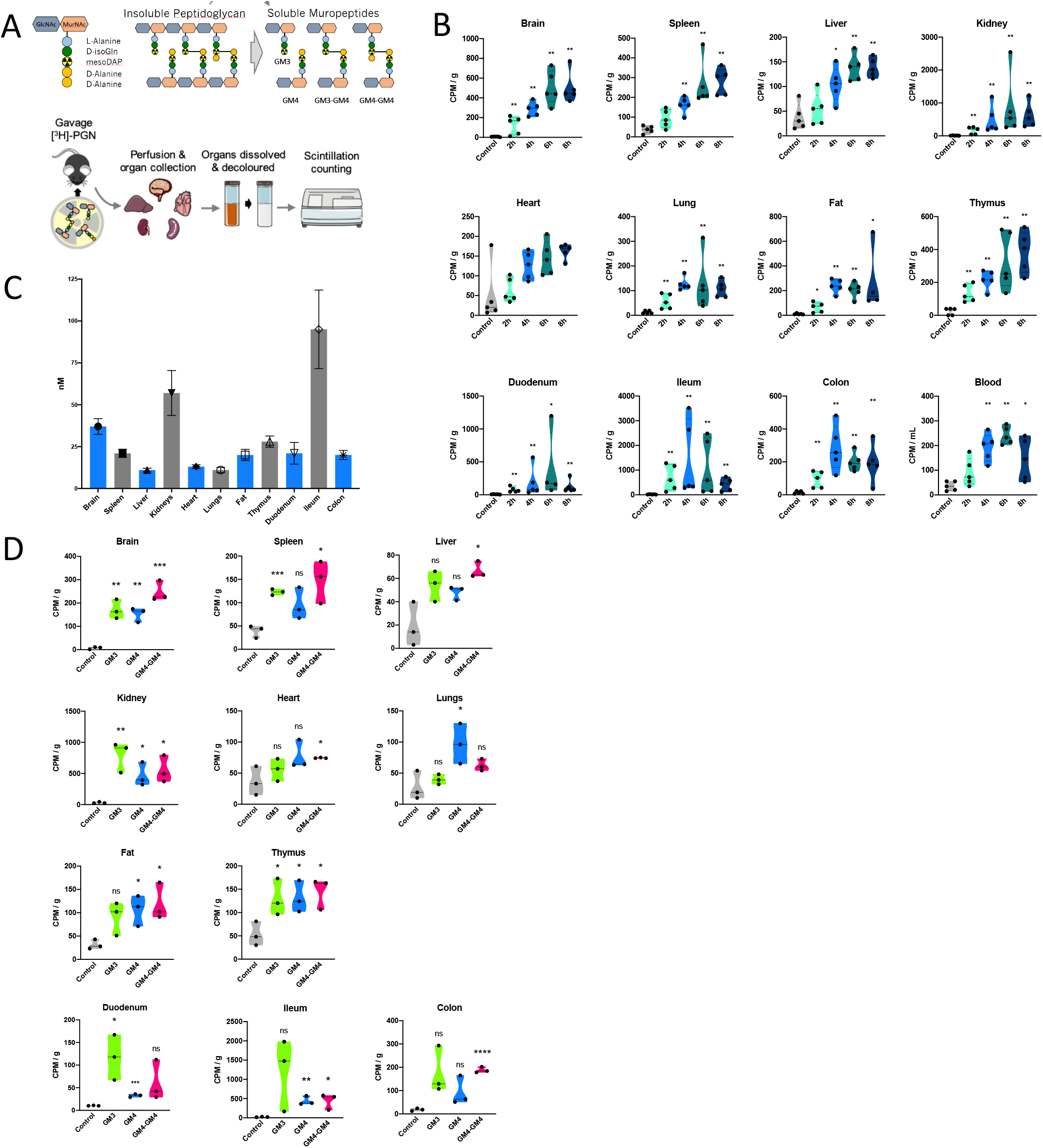
The kinetics of [3H]-PGN biodistribution following administration by gavage in mice. A) Schematics summary of Radiolabeling methodology, indicating the position of the 3H-labelled mesoDAP, and radiotracking strategy. Mice (female C57BL/6j, 8-12 weeks) were gavaged with [3H]-PGN. At the desired timepoint, mice were perfused to clear transiently circulating [3H]-PGN from the organs. The tissues to be analyzed were dissolved, then decolored by peroxide bleaching to reduce quenching effects, prior to scintillation counting. B) [^3^H]-PGN measured by scintillation counting of dissolved, decolored organs and blood between 2h and 8 h post-gavage. Measured CPM values are normalized per g tissue weight, or per mL blood. Pairwise comparison with control performed using the Mann–Whitney U test. \**p* ≤ 0.05; ** p ≤ 0.0050. Welch’s ANOVA comparing time-point groups, excluding control: Brain *p* = 0.0045; Spleen *p* = 0.0050; Liver *p* = 0.0068; Kidney *p* = 0.087; Heart *p* = 0.0029; Lung *p* = 0.0331; Fat *p* = 0.0043; Thymus *p* = 0.0165; Duodenum *p* = 0.4681; Ileum *p* = 0.3505; Colon *p* = 0.0375; Blood *p* = 0.0082. C) Estimated concentration of [^3^H]-PGN in each organ. Concentration of [3H]-PGN per CPM was determined by HPLC analysis of a predetermined CPM value of [3H]-PGN. The total peak area was calculated and the value in mg determined relative to an MDP standard curve. D) The biodistribution of [^3^H]-GM3, [^3^H]-GM4 and [^3^H]-GM4-GM4, administered *per os* in SPF mice. Scintillation counting was performed on dissolved, decolored organs. Results are normalized to CPM per g of tissue. Pairwise comparison with control performed using unpaired t-test. * *p* ≤ 0.05; ** p ≤ 0.0050; *** p ≤ 0.0005; p < 0.0001. Welch’s ANOVA comparing muropeptides groups, excluding control: Brain *p* = 0.0582; Spleen *p* = 0.2711; Liver *p* = 0.1508; Kidney *p* = 0.2478; Heart *p* = 0.3525; Lung *p* = 0.1022; Fat *p* = 0.6536; Thymus *p* = 0.8542; Duodenum *p* = 0.1234; Ileum *p* = 0.3295; Colon *p* = 0.2227.

At two-hours post-gavage with [^3^H]-PGN, radioactivity was detected differentially across the host organs **(Fig. 1B and fig. S2A)**. Normalized by tissue weight, the highest levels of exogenously administered peptidoglycan were observed in the brain, spleen, kidneys, thymus and fat, whereas the lowest levels were observed in the liver, heart and lungs (for absolute counts per organ, see **Fig. S2A**). This relationship did not change over time. The presence of radioactivity in the organs accumulated, reaching maximal values at 6-8 hours post-gavage. Similarly, radioactivity in the blood peaked at 6h, then declined slightly at 8h. Values in the intestinal tract showed the greatest variability over time, likely reflecting the local presence of peptidoglycan undergoing peristaltic transit through the intestinal lumen, or coprophagic behavior by some mice.

To quantify exogenously administered peptidoglycan in the tissues, the ratio of ^3^H-*meso*DAP per unit weight of [^3^H]-PGN was determined by scintillation counting, combined with reversed-phase HPLC quantification of [^3^H]-PGN muropeptides against a standard curve of muramyldipeptide. Based on the values measured in the organs between 6h and 8h, we determined that the peptidoglycan accumulated in the order of 5-55 nanograms per gram of tissue, and 20 ng/mL in the blood **(Fig. 1C)**. Our data are consistent with reported levels of serum peptidoglycan in SPF mice, measured by indirect, competitive enzyme-linked immunosorbent assay, which also fall in the ng/mL range (180-300 ng/mL), demonstrating the physiological relevance of our experimental approach^8^.

The selectivity of the gut epithelial barrier toward peptidoglycan fragments is a critical parameter that could determine the nature of the peptidoglycan moieties found systemically. A variety of mechanisms have been proposed for epithelial transit of peptidoglycan fragments, ranging from non-specific mechanisms such as paracellular transport and carrier-mediated mechanisms, to highly specific mechanisms targeting a restricted set of monomeric muropeptides, such as the SLC15 family peptide transporters, suggested to transport muramyl dipeptides and tripeptides, reviewed in^11,12^. To gain insight into the selectivity of the intestinal epithelial barrier towards different muropeptides, we gavaged mice with 50,000 counts per minute (cpm) of individual, purified ^3^H-labelled muropeptides possessing a peptide stem length of three, four or eight amino acids; the monomers [^3^H]-GM3, [^3^H]-GM4, or the dimer [^3^H]-GM4-GM4. The systemic presence of the ^3^H-labelled muropeptides was assessed 4h post-gavage **(Fig. 1D, Fig. S2B)**. In all cases, ^3^H was detected in approximately equal proportions across the distinct organs, demonstrating that muropeptide size is not a limiting factor for intestinal absorption and dissemination process.

To assess the possibility that [^3^H]-PGN fragments are degraded in the stomach and gut, resulting in the liberation and absorption of mainly free [^3^H]-*meso*DAP, we assessed whether [^3^H]-*meso*DAP and [^3^H]-PGN displayed distinct kinetics of absorption and dissemination **(Fig. 2A, fig. S3A)**. Gavage with [^3^H]-*meso*DAP did not recapitulate the kinetics or distribution profile observed for [^3^H]-PGN. Notably, [^3^H]-*meso*DAP reached maximal values in the kidney at 2h post-gavage, followed by rapid clearance (**Fig. 2A, fig. S3A)**. Abundant accumulation in the fat peaked between 2h and 6h, and then declined. Uptake in the spleen and liver reached their maximum by 2h and remained steady for the duration of the experiment. Only comparably low levels of [^3^H]-*meso*DAP accumulated in the brain, differing from the case of [^3^H]-PGN where relative to the other organs, the brain was a major reservoir. The striking differences between the dynamics of [^3^H]-*meso*DAP and [^3^H]-PGN indicate that peptidoglycan does not undergo acute catabolism in the gut leading to absorption of mainly free *meso*DAP.

**Fig. 2.**
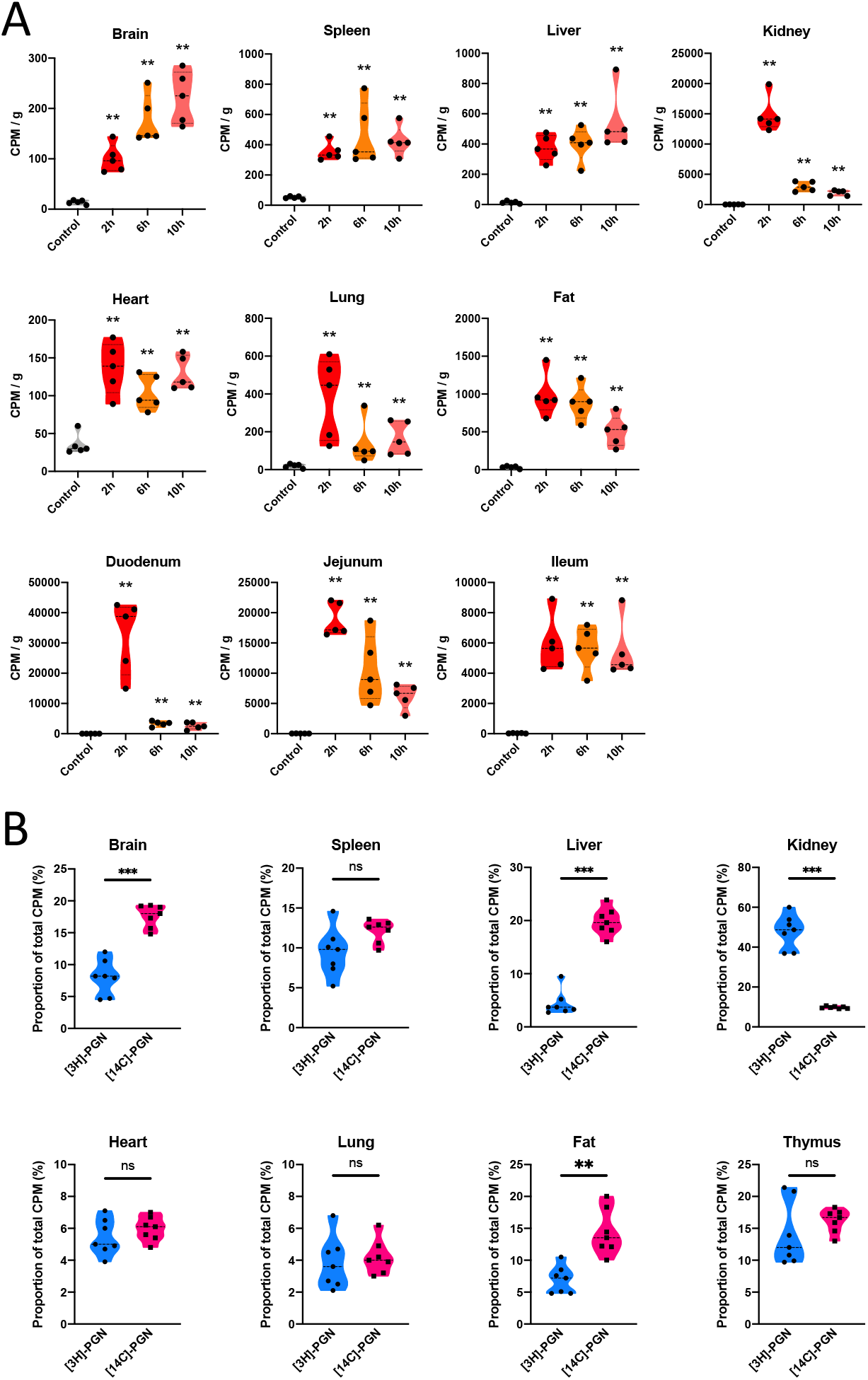
Effect of labelling strategy parameters on biodistribution of peptidoglycan. A) Biodistribution of [^3^H]-mesoDAP amino acid administered to mice *per os*. [^3^H]-mesoDAP biodistribution was measured by scintillation counting of dissolved, decolored organs, 2h, 6h and 10 h post-gavage. The measured CPM values are normalized per g tissue weight. B) Biodistribution of [^3^H]-PGN versus [^14^C]-PGN administered to mice by gavage. The relative abundance of [^3^H]-mesoDAP or [^14^C]-GlcNAc labelled peptidoglycan detected for each organ or tissue. Data are presented as the proportion of the sum of all [3H] or [14C] CPM measurements in the organs per mouse. Pairwise comparison with control performed using the Mann–Whitney U test. \*\**p* ≤ 0.005; \*\*\**p* ≤ 0.0005. Welch’s ANOVA comparing timepoint groups, excluding control, Brain *p* = 0.0052; Spleen *p* = 0.3334; Liver *p* = 0.3307; Kidney *p* = 0.0002; Heart *p* = 0.1921; Lung *p* = 0.1624; Fat *p* = 0.0323; Duodenum *p* = 0.0040; Jejunum *p* = 0.0003; Ileum *p* = 0.9333.

PGLYRP-2 serum amidase cleaves muropeptides larger than MDP, generating peptides that are preferentially cleared via urinary excretion^13^. To gain insight into the integrity of disseminated peptidoglycan fragments, we compared the distribution profile of orally administered peptidoglycan when the glycan chain was radiolabeled versus the peptide stem. Mice were gavaged with 400,000 cpm of peptidoglycan labelled with [^14^C]-*N*-acetyl-D-glucosamine ([^14^C]-PGN) or [^3^H]-PGN. Since the two radiolabels have different activities and levels of incorporation, their dissemination profiles cannot be compared directly. We therefore plotted the proportion of ^3^H or ^14^C in each organ, relative to the total amount of ^3^H or ^14^C radioactivity measured systemically, excluding the intestinal tract **(Fig. 2B, fig. S3B and C)**. For most of the organs, the ratio of [^3^H]-PGN to [^14^C]-PGN was approximately 1:1, suggesting that these are intact peptidoglycan fragments that escaped serum amidase activity. The brain and fat showed small deviations, with the proportion of [^14^C]-PGN approximately double that of [^3^H]-PGN, suggesting separate accumulation of glycan versus intact muropeptide or peptide fractions. The kidneys and liver demonstrated notable deviations, with [^3^H]-PGN to [^14^C]-PGN ratios of 5:1 and 1:4, respectively. Our data are consistent with a report that peptides are preferentially cleared via the kidney (urinary excretion), whereas free disaccharide is preferentially directed to the liver^14^.

We next asked whether the systemic distribution of [^3^H]-PGN is altered when the intestinal barrier is bypassed completely. We administered 40,000 cpm of [^3^H]-PGN intravenously, an amount that exceeds the maximal values detected systemically in gavage experiments, and measured accumulation in organs at 1h, 4h and 8h post-injection (p.i.). Surprisingly, with the exception of the kidney, [^3^H]-PGN was detected at low levels across the organs **(Fig. 3A and Fig. S4A)**. Peak accumulation of [^3^H]-PGN in the kidney occurred 1h p.i., followed by rapid clearance, such that close-to-background levels were reached by 4h p.i., suggesting rapid clearance of this dose of [^3^H]-PGN via urinary excretion. Only when the system was forced by intravenous administration of 400,000 cpm of [^3^H]-PGN, the same dose administered in gavage experiments, did we observe uptake in the organs at 1h p.i. **(Fig. 3B and Fig. S4B)**. When compared to [^3^H]-PGN biodistribution at 4h post-gavage, intravenously administered [^3^H]-PGN was detected at similar or slightly elevated levels in the spleen, liver, kidney, lungs, fat and thymus, and at a reduced level in the heart. Strikingly, intravenously administered [^3^H]-PGN was barely detectable in the brain. We next administered 400,000 cpm of [^3^H]-PGN intraperitoneally. Maximal uptake to the kidneys, spleen, liver, fat and thymus was observed at the earliest time point (2h p.i.), similar to the kinetic profile obtained by intravenous administration of [^3^H]-PGN **(Fig. 3C and Fig. S4C)**. Radioactivity in kidneys and fat was lower at 6h p.i., indicating clearance, whereas [^3^H]-PGN levels were sustained in the spleen and liver at 6h p.i. [^3^H]-PGN was not detected in the heart, lungs, or intestinal tract. As with intravenously administered [^3^H]-PGN, intraperitoneally administered [^3^H]-PGN was not detected in the brain. Together, these data suggest that entry to the brain is highly restricted when [^3^H]-PGN is administered directly into the host system, bypassing the gut, whilst absorption across the intestinal barrier facilitates subsequent trafficking to the brain.

**Fig. 3.**
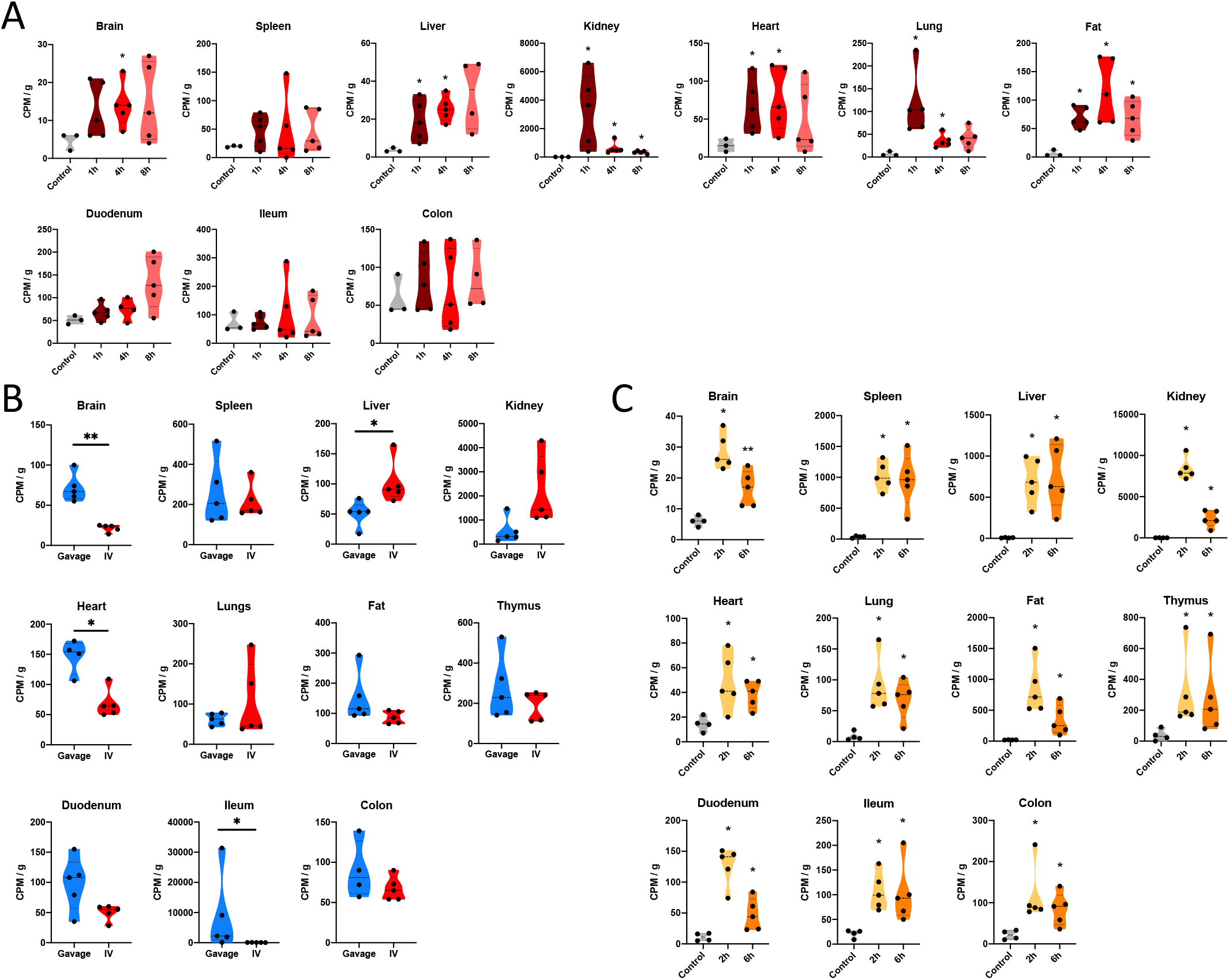
Dose-dependent biodistribution of [^3^H]-PGN administered intravenously or intraperitoneally. A) Mice were administered 40,000 CPM of [3H]-PGN intravenously and biodistribution to organs and tissues measured at 1h, 4h, 8h and 24h post-injection. Welch’s ANOVA comparing time-point groups, excluding control: Brain *p* = 0.9236; Spleen *p* = 0.9972; Liver *p* = 0.4364; Kidney *p* = 0.0842; Heart *p* = 0.6337; Lung *p* = 0.1320; Fat *p* = 0.2897; Duodenum *p* = 0.1497; Ileum *p* = 0.8240; Colon *p* = 0.9038. B) Mice were administered 400,000 CPM of [3H]-PGN intravenously (two injections of 200,000 cpm) 30 minutes apart, or 400,000 CPM of [3H]-PGN by gavage. Organs were harvested at 1h after the first injection, or 4h after gavage. Scintillation counting was performed on the dissolved, decolored organs. C) Biodistribution of [^3^H]-PGN administered intraperitoneally. Mice were injected intraperitoneally with 400,000 CPM of [^3^H]-PGN and by scintillation counting performed on the dissolved, decolored organs harvested at 2h and 6h post-gavage. The measured CPM values are normalized per g tissue weight, Data are normalized as CPM values per g tissue weight. Pairwise comparison with control performed using the Mann–Whitney U test. \**p* ≤ 0.05; ** *p* ≤ 0.005. C) Pairwise comparison with between time-points performed using the Mann–Whitney U test, Brain *p* = 0.0159; Spleen *p* = 0.8413; Liver *p* = 0.8413; Kidney *p* = 0.0079; Heart *p* = 0.7222; Lung *p* = 0.5952; Fat *p* = 0.0317; Duodenum *p* = 0.0159; Ileum *p* = 0.6905; Colon *p* = 0.8413.

Having identified that the intestinal tract functions as a gateway favoring the accumulation of peptidoglycan through the host system, we investigated the initial step of peptidoglycan translocation across the small intestinal epithelial barrier using confocal microscopy. Fluorescent conjugates of MDP (MDP-rho), or *E. coli* peptidoglycan (Fluo-PGN) were injected into ileal ligatures of anaesthetized mice. After 20 minutes of incubation, the tissue was washed, fixed and stained for imaging by confocal microscopy. We observed transcellular uptake of MDP-rho and Fluo-PGN in the ileal villus epithelium **(Fig.4A and B, fig. S5)**. Goblet cells (identified by peripheral cell mucus-staining with wheat germ agglutinin, cellular morphology and basal positioning of the nucleus) displayed a prominent affinity for MDP-rho and Fluo-PGN, with fluorescent peptidoglycan observed as a ring around the apical portion of the cell, excluded from the regions containing mucus, and more diffuse in the basal cytoplasm **(Fig. 4A and B, fig. S5)**. The intensity of fluorescence in these cells was variable, perhaps reflecting the local concentration of peptidoglycan or dynamics of uptake by these cells. We did not observe PGN translocation in enteroendocrine cells, tuft cells, or in microfold cells **(Fig. S5C)**.

**Fig. 4.**
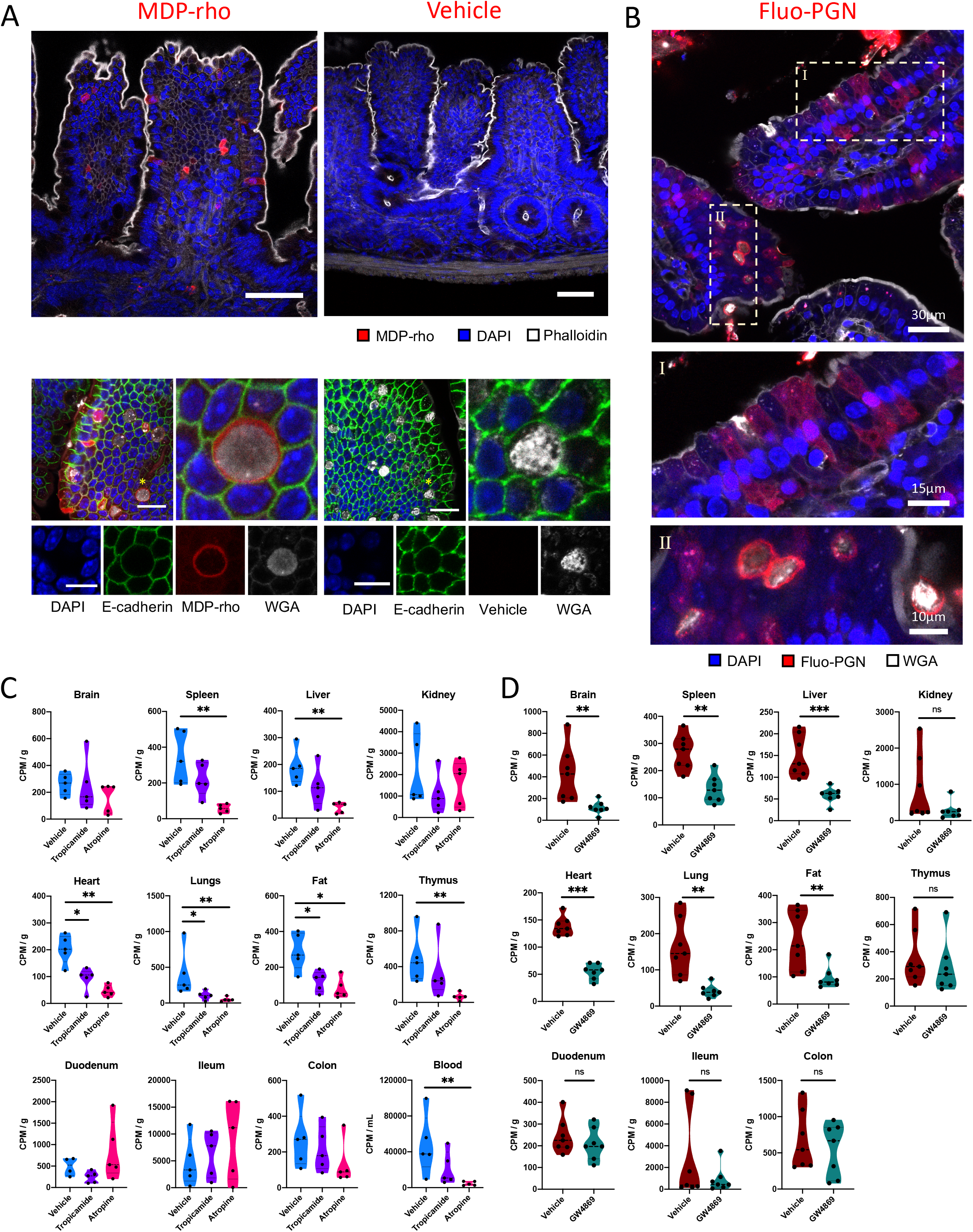
Peptidoglycan uptake and dissemination by intestinal epithelial cells. A) Upper panel; MDP-rho internalization was observed in a subset of epithelial cells in the ileal villi. Lower panel; internalization of MDPrho in goblet cells (WGA^+^). No fluorescence was observed in the villus epithelium including the goblet cells, in the rhodamine channel of vehicle treated controls. B) Internalization of PGN-AF647 in the ileum of SPF mice by a subset of epithelial cells, which includes goblet cells (WGA^+^; panel I) and other cell types (WGA^-^, panel II). C) The biodistribution of [^3^H]-PGN administered *per os* in SPF mice, is suppressed by parasympathetic inhibition. SPF mice were administered tropicamide, atropine or vehicle control prior to gavage with [^3^H]-PGN. Scintillation counting was performed on dissolved, decolored organs. D) [^3^H]-PGN biodistribution from the gut is suppressed by GW4869 treatment. Results are normalized to CPM per g of tissue. Pairwise comparison to vehicle control performed using the Mann–Whitney U test. \**p* ≤ 0.05; ** *p* ≤ 0.005

Microscopy analysis indicated that peptidoglycan transits the epithelial barrier via active transepithelial processes, and prominent uptake by Goblet-cells. Goblet-cell associated antigen passages (GAPs) translocate luminal antigens to underlying antigen presenting cells, facilitating small intestinal tolerogenesis^15^, a process regulated by muscarinic acetylcholine receptor (mAchR)-4 activation^16^. Expression of all five mAchR subtypes is reported in the mouse small intestine epithelia^17–19^. We asked whether mAchR-mediated pathways regulate the translocation and dissemination of peptidoglycan. We performed *per os* [^3^H]-PGN biodistribution analysis in mice treated with the parasympatholytic agents tropicamide (selective antagonist of mAchR-4) or atropine (pan-mAchR antagonist). Pan-mAchR antagonism with atropine led to significant inhibition of [^3^H]-PGN biodistribution **(Fig. 4C and fig. S6A)**. Atropine treatment significantly reduced the levels of [^3^H]-PGN circulating in the blood, suggesting that mAchR inhibition restricts peptidoglycan absorption at the level of the intestine. In the case of tropicamide-treatment targeting peptidoglycan uptake via goblet cells, we observed a trend of reduced distribution to the organs and blood, although for the majority of organs this reduction did not meet the significance cut-off of *p* ≤ 0.05. Our data suggest a partial role for GAPs in the systemic dissemination of PGN.

*In vitro* studies suggest that peptidoglycan is exocytosed by intestinal epithelial cells via exosomes, a potentially mechanism underlying the distinct organ tropism of peptidoglycan arriving from the gut^20^. To assess the possibility of peptidoglycan dissemination via exosomes, we inhibited exosome generation by administering neutral sphingomyelinase inhibitor GW4869 to mice, followed by gavage with [^3^H]-PGN^21^. Radioactivity presence was diminished in almost all organs tested following GW4869 treatment, but not in the gut, suggesting that blockade of exosome confines peptidoglycan in the gut tissue **(Fig. 4D and fig. S6B)**. Although we cannot rule out indirect effects, our data support a significant role for exosomes in the gut peptidoglycan dissemination pathway.

In mice lacking a microbiota (germ-free), the gut is considered to be leaky, as microbial colonization triggers tightening of the intraepithelial tight junctions, augmenting epithelial paraselectivity. To explore the connection between the gut microbiota, intestinal permeability and systemic biodistribution of peptidoglycan, we compared the dissemination of [^3^H]-PGN administered by gavage to SPF mice, germ-free mice, and formerly germ-free mice, conventionalized with SPF microbiota. Remarkably, peptidoglycan biodistribution was suppressed in germ-free mice, with low levels of [^3^H]-PGN detected across the distinct organs **(Fig. 5A and fig. S7A)**. By comparison, peptidoglycan biodistribution was restored to SPF levels upon conventionalization **(Fig. 5A, fig. S7A and B)**. The observed effect was not due to impaired intestinal motility in germ-free mice, since tracking of [^3^H]-PGN administered by gavage showed that gut peptidoglycan reached the jejunum and cecum at 2 hours and 8 hours respectively with low presence in organs **(Fig. S7C)**. Epithelial cell uptake of MDP-rho was observed in SPF, but not germ-free mice, suggesting that the passage of peptidoglycan was restricted at the level of the intestinal epithelial barrier **(Fig. 5B)**. To investigate whether the uptake of intestinal peptidoglycan observed in SPF mice requires the sustained presence of the microbiota or is due to an irreversible physiological alternation triggered by microbial colonization, we depleted the microbiota of adult SPF mice by administering a cocktail of antibiotics in the drinking water for 7 days prior to gavage with [^3^H]-PGN. Antibiotic depletion of the microbiota suppressed the systemic biodistribution of gut peptidoglycan **(Fig. 5C and fig. S7D)**, demonstrating that absorption of gut peptidoglycan is malleable to changes in microbial colonization status.

**Fig. 5.**
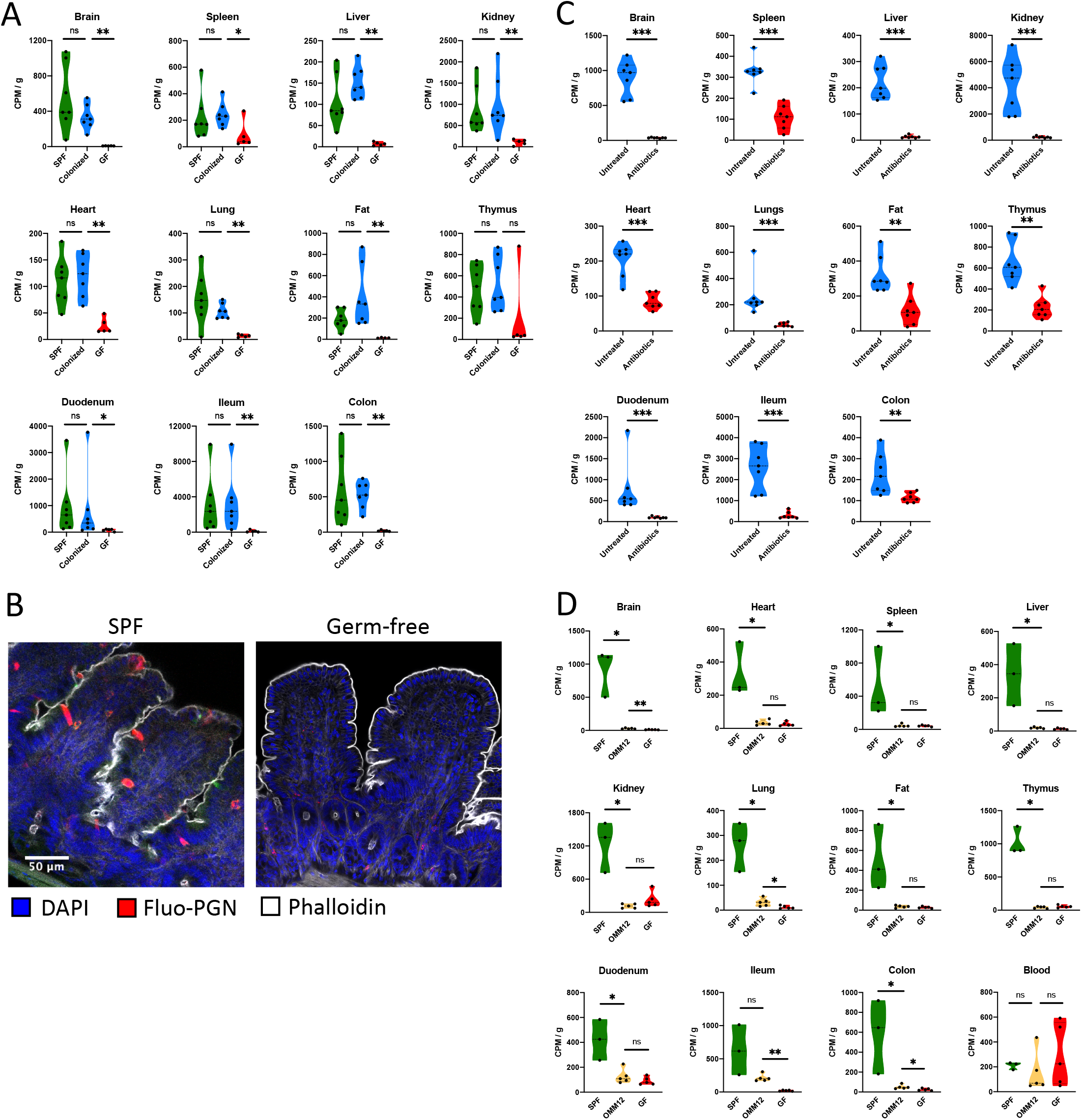
Microbiota colonization is required for efficient PGN biodistribution. A) The biodistribution of [^3^H]-PGN in germ-free (GF) mice, specific pathogen free (SPF) mice and colonized mice (previously GF mice co-housed with SPF for 3 weeks). B) Epithelial cell uptake of MDP-rho observed in the ileal villi of SPF mice is absent in germ-free mice. C) Biodistribution of [3H]-PGN in SPF mice treated with a broad-spectrum antibiotic cocktail (ATX). D) The biodistribution of [^3^H]-PGN in OMM12 mice, compared with SPF and germ-free mice. Scintillation counting was performed on dissolved, decolored organs. Results are normalized to CPM per g of tissue. Pairwise comparison to GF condition performed using the Mann–Whitney U test. * *p* ≤ 0.05; ** *p* ≤ 0.005; *** *p* ≤ 0.0005

Finally, we assessed [^3^H]-PGN absorption and dissemination in Oligo-Mouse-Microbiota (OMM^12^) mice, which are colonized by a functional synthetic microbiota of twelve bacterial species representing the five major Phyla naturally abundant in the mouse gut^22^. We observed that OMM^12^ microbiota colonization was not sufficient to restore peptidoglycan absorption and dissemination pathways, having a profile similar to that of germ-free mice **(Fig. 5D and fig. S7E)**. Taking these data together, we conclude that the presence and composition of the gut microbiota are critical determinants to activate and sustain the systemic biodistribution of peptidoglycan.

## Discussion

In this study, we explored fundamental parameters governing the biodistribution of peptidoglycan originating in the mammalian host gut. Our study highlights that the kinetics and tropism of systemically distributed peptidoglycan depend on the route of administration. Orally administered peptidoglycan gradually disseminated to major organs over 6 h. Direct intravenous delivery of peptidoglycan was detected primarily in the kidneys where the peptidoglycan presence was high at 1h, then declined rapidly, indicating that intravenously administered peptidoglycan is mainly excreted via the urine. Consistent with our data, previous studies tracking intravenously administered peptidoglycan moieties showed that up to 80% of 14C labelled muropeptide monomers were excreted within the first hour^14,23^. Fewer studies have explored the fate of peptidoglycan entering the host system via the gut. Valinger *et al*^24^, showed that 14C-labelled peptidoglycan disaccharide pentapeptide monomer administered by oral gavage gradually accumulated in organs over several hours. Only 15% of radioactivity was excreted in the urine, whilst up to 30% of radioactivity was exhaled as CO^2^ within 48h. Host metabolic pathways that degrade peptidoglycan to CO^2^ or other products have not been identified. Serum amidase PGLYRP-2 and lysozyme remain the only host enzymes known to directly degrade peptidoglycan in mammals. Subsequently, targeted mass spectrometry experiments performed on plasma or serum of different mammalian hosts have detected only a small number of anticipated masses associated with classical NOD1 and NOD2 ligands^8,9^. In this study, we demonstrated that when mice were gavaged with radiolabeled muropeptides with a peptide stem length of three, four or eight amino acids, radioactivity was detected in approximately equal proportions in the organs, and in blood. Thus, muropeptides of different sizes are equally capable of acting as a source of the peptidoglycan moieties disseminated systemically. Our untargeted approach followed radioactivity from the *meso*DAP moiety of peptidoglycan, and thus potentially detects uncharacterized products of host peptidoglycan metabolism derived from the *meso*DAP component. The host is increasingly found to respond to non-classical peptidoglycan derivates, independent of NOD1 and NOD2 receptors^25,26^, and even subtle structural differences between naturally occurring muramyldipeptides and muramyltripeptides are reported to induce different innate immune response profiles upon PRR activation^27^. The importance of enzymatic processing of peptidoglycan in the intestine is yet to be fully explored, and may influence tropism by enriching the pool of moieties that are substrates for receptors or transporters at epithelial and endothelial barriers. Lysozyme activity was already shown to be critical to the systemic availability of NOD1 ligands^28^.Thus, our data further highlights the need to better understand the true scope of peptidoglycan metabolism by the host.

A striking result was that peptidoglycan was only detected in the brain when administered by gavage. Intravenously administered peptidoglycan was not detected in the brain, and absence from the brain was even more striking when administered intraperitoneally, since a higher relative abundance of peptidoglycan was observed in the spleen, liver, kidney and fat. Growing evidence suggests a gut-brain axis connecting microbiota peptidoglycan translocation to effects on the brain^10,29,30^. Our data indicate that for studies of peptidoglycan effects on the brain, the route of delivery is a critical factor. Intestinal transit facilitates entry to the brain, perhaps by enabling crossing of the blood-brain barrier (BBB), whilst the systemic presence of peptidoglycan does not guarantee presence in the brain. Peptidoglycan administered by gavage disseminated to the brain with the same kinetic profile as for other organs, suggesting a shared pathway of translocation. An open question is whether gut peptidoglycan is delivered to organs via a vehicle such as a carrier protein, vesicles or cells that could protect against serum amidase activity and facilitate transit of barriers such as the BBB. It has been proposed that macrophages, dendritic cells and neutrophils could perform this function^29^, whilst our data add to evidence for a role of exosomes in peptidoglycan trafficking^20^.

In this study, we demonstrated that the pathway facilitating intestinal translocation of peptidoglycan is induced by the gut microbiota. Peptidoglycan absorption and dissemination in germ-free mice occurred at extremely low levels compared with SPF mice. This basal level of absorption in germ-free mice is nevertheless sufficient to facilitate peptidoglycan-mediated effects in the gut and systemically^3,6^. The germ-free gut is considered leaky due to the observation, that microbiota colonization, or microbial metabolites, induce fortification of the intestinal barrier junction integrity^31–34^. Our data suggest that in the absence of epithelial junction tightening, barrier integrity is sufficient to restrict the passive permeation of peptidoglycan fragments. This finding is consistent with a previous report that the germ-free mouse colon is less permeable to MDP and lipopolysaccharide than that of conventional mice^35^. In germ-free mice the peptidoglycan uptake pathway was induced by colonization with conventional microbiota, but not by colonization with the OMM^12^ synthetic microbiota representing five major phyla of the conventional mouse gut. Conversely, peptidoglycan absorption was suppressed in adult SPF mice by antibiotic depletion of the microbiota. Our data suggest that specific members of the microbiota directly or indirectly induce and sustain the pathway of gut peptidoglycan absorption. Fluorescent muropeptide uptake was observed in gut epithelia of SPF mice, but not germ-free mice, indicating that peptidoglycan absorption is restricted at the level of the epithelial interface with the gut lumen.

Despite the robust uptake of fluorescently labelled PGN by goblet cells, we found that GAPs play a partial role in peptidoglycan systemic dissemination. Goblet cells facilitate delivery of luminal antigen to innate immune regulatory cell populations such as CD103+ dendritic cells (DCs), to promote lamina propria immune cell tolerance to dietary antigens^15,17^.

*Lactobacillus salivarius* Ls33 peptidoglycan was found to protect mice against experimental colitis via a CD103+ DC dependent mechanism^36^, supporting a hypothesis that goblet cell absorption of peptidoglycan is primarily an immune-sampling function. M-cells, another specialist cell class involved in immune surveillance, are proposed to absorb nanomineral encapsulated peptidoglycan for presentation to antigen-presenting cell sentinels^37^. We did not observe peptidoglycan absorption by M-cells, however our methodology (injection of fluorescent muropeptide conjugate into ileal ligatures over a short time frame) may not favor detection of nanomineral encapsulated peptidoglycan. Although we cannot exclude GAP-independent trafficking of peptidoglycan by Goblet-cells, our data support a hypothesis of functional separation between epithelial cells involved in luminal peptidoglycan sampling for immune surveillance, and those that coordinate the steady-state dissemination of peptidoglycan.

This study highlights the importance of the gut-axis for the systemic tropism of peptidoglycan and reveals major processes regulating the dissemination of peptidoglycan from the gut. By exploring such parameters, we will better understand the peptidoglycan-driven mechanisms through which gut microbiota mediate steady-state processes in health, and how their dysbiosis drives pathological effects.

## Acknowledgments

We acknowledge the UTechS PBI a member of the France–BioImaging infrastructure network supported by the French National Research Agency (ANR-10–INSB– 04, Investments for the future) for microscope usage and assistance.

## Author contributions

Conceptualization PB, IB and RW; methodology development: radiotracking PB and RW, radio-HPLC, LC-MS AR and RW Investigation PB, RW, AR, JS, OD; Formal analysis PB and RW; Resources: IB; Writing - Original Draft: RW, PB, IB Review & Editing: PB, IB, RW Visualization: PB and RW; Supervision: IB, ML and RW; Project administration: PB and RW; Funding acquisition: IB.

## Data and materials availability

All data is available in the main text or the supplementary materials.

## Materials and Methods

### Radiolabeling and purification of peptidoglycan

Radiolabeling was performed in the *E. coli* FB8-LysA strain which is unable to convert meso-diaminopimelic acid (mesoDAP) to lysine by decarboxylation^38^. Therefore, mesoDAP added to the growth media is incorporated specifically into the peptidoglycan layer. *E. coli* FB8-LysA was cultured on LB Miller agar (BD Difco) containing 25 mg/ml kanamycin (Merck). Several colonies were inoculated into 50 mL of LB medium plus 25 mg/ml kanamycin and incubated at 37°C. At approximately OD600 1.0, the pre-culture was used to inoculate (1:100) 1L of prewarmed M9 minimal media supplemented with 100 μg/ml threonine, methionine and lysine (Merck). For 3H-mesoDAP labelling, 50 μCi/L of 3H-meso diaminopimelic acid (3H-mesoDAP; Moraveck Inc.) was added. For 14C-GlcNAc labelling, culture was performed in M9 media using 100 μM GlcNAc (Merck) as the carbon source, and spiked with 10 μCi/L of ^14^C-*N*-acetylglucosamine (^14^C-GlcNAc; ARC). Cultures were incubated overnight at 37°C with aeration. Final OD600 was approximately 2.0. Bacteria were harvested by centrifugation at 4000 xg, resuspension in a small volume of cold H_2_O, and dropped into 20 mL of 4% SDS in a boiling bain-marie. After 1 hr of boiling with vigorous agitation, the suspension was cooled to room temperature, and centrifugated at 13,000 xg. The supernatant was discarded, and the pellet resuspended with 20 mL of H_2_O. The radiolabelled cell wall material was washed by centrifugation and resuspension of the pellet with H_2_O until the presence of SDS in the supernatant could no longer be detected using the method of Hayashi^39^. The pellet was then resuspended in 4 mL of 50 mM Tris pH 7.5 (Merck), and incubated at 37°C with 100 μg/mL alpha-amylase (Merck) for 2 h, followed by 2h incubation with RNase, DNase and MgCl2, then incubated overnight with 100 μg/mL 3x crystallized trypsin (Worthing Biochemical Corporation) and CaCl2. The pellet was then incubated for 15 min at 100 °C, then washed once with H_2_0. Radiolabelled peptidoglycan was stored at −20 °C. For gavage of mice, peptidoglycan was digested overnight with 100 U/mg of mutanolysin (from *Streptomyces globisporus* ATCC 21553, Merck) in 12.5 mM sodium phosphate pH 5.6 (Merck) at 37°C.

### Animals

Mice were housed at the Institut Pasteur animal facilities under specific pathogen-free conditions. Female mice between 8 and 12 weeks of age were used for all experiments. C57BL/6J mice were purchase from Charles River Laboratories. NOD1/2 dKO mice, germ-free mice and OMM12 mice were bred and maintained at Institut Pasteur facilities.

### Ethics statement

Animal experiments were performed in accordance with Directive 2010/63/EU of the European Parliament and the French regulation for the protection of laboratory animals decree of February 1, 2013. The project number APAFIS #8551 was approved by the Institut Pasteur ethical committee for animal experimentation (Comité d’Ethique en Expérimentation Animale CETEA registry number #89) and authorized by the Ministère de l’Enseignement HSupérieur, de la Recherche et de l’Innovation (MESRI).

### Radioactive peptidoglycan tracking

Mice were gavaged with a 200 μL volume containing approximately 400,000 cpm of labelled peptidoglycan. Approximately 30 minutes before the specified time point, profound anaesthesia was induced in mice by intraperitoneal administration of 100 mg/kg Imalgene1000 (Boehringer-Ingelheim) and 8 mg/kg of Rompun 2% (Bayer). To avoid the detection of radioactivity transiently present in the circulation, blood was cleared from the organs by transcardial perfusion with 40 mL of PBS (Lonza Pharma & Biotech) by syringe with 26G needle, with section of the inferior vena cava. Blood collection was performed immediately upon section of the vena cava. Organs were removed into 2 mL microfuge tubes. For small intestine samples, approximately 8 cm of tissue adjacent to the stomach (“Duodeum”), and 8 cm of tissue adjacent to the cecum (“Ileum”) were collected. Approximately 6 cm of colon as collected. The gut pieces were opened longitudinally, then the gut contents removed by vigorous washing three-times in PBS. The gut tissue was dried on tissue paper then transferred to 2 mL microfuge tubes. For blood, collection tubes contained 10 μL of 0.5 M EDTA pH 8.0 (Lonza Pharma & Biotech). The weights of all organs and tissues were recorded. If further processing was not performed immediately, the samples were stored at −20 °C.

### Drug treatments

Parasympatholytic reagents: 100x stocks of tropicamide (Merck) in ethanol (5.5 mg/mL) or atropine sulfate (Merck) in ethanol (6 mg/mL) were prepared and stored at 4°C. On the day of injection, reagents were diluted 1:100 in PBS. Tropicamide (550 μg/kg) or atropine sulfate (600 μg/kg) were injected intraperitoneally in a 200 μL volume, 20 min prior to administration of [3H]-PGN. Control mice received vehicle only.

Exosome inhibition: Five mg/mL of GW4869 (N,N’-Bis[4-(4,5-dihydro-1H-imidazol-2-yl)phenyl]-3,3’-p-phenylene-bis-acrylamide dihydrochloride; Merck) solution was diluted in sterile PBS to a final concentration of 0.25 mg/mL. Two-hundred microlitres of GW4869 solution was administered intraperitoneally 16h and 2h prior to gavage with [3H]-PGN. Control mice received vehicle only.

Antibiotic treatment: intestinal microbiota of SPF mice was depleted by administration of antibiotic cocktail in the drinking water of mice for 21 days. Ampicillin (1 mg/mL, Merck), Streptomycin (5 mg/mL, Euromedex), Colistin sulfate (1 mg/mL, Merck) and Vancomycin (0.25 mg/mL, Merck) were prepared in water and filtered through 0.22 μm sterile filters (Corning) and protected from light with aluminium foil. Fresh solution was changed every 3 days. Microbiota depletion was regularly checked by spotting 10 μl serial dilutions of fecal homogenate in PBS onto trypticase soy agar (BD) enriched with 5% defibrinated horse blood (Thermo Fisher) and incubation at 37°C under aerobic and anaerobic atmosphere. Colony forming units were counted after 24h and 48h of incubation.

### Scintillation counting

Organs, tissues or blood (200 μL) were transferred into glass vials, then dissolved by addition of 2mL (3 mL for the liver samples) of Solvable (PerkinElmer) to each vial. Samples were incubated overnight at 60 °C. After cooling to room temperature, the solutions were rendered transparent by addition of 100 μL of 0.5 M EDTA pH 8, followed by 2x 200 μL of 30% hydrogen peroxide (Millipore). Samples were incubated for 30 min at 60 °C. After cooling again to room temperature, the solutions were poured into 20 ml HDPE scintillation vials with urea cap containing a polyethylene cone (Duran Wheaton Kimble), followed by addition of 10 mL of Ultima Gold LLT scintillation cocktail (Perkin Elmer). The samples were equilibrated in the dark at room temperature for 4 hours. Scintillation counting was performed using a Tri-Carb 3110 TR Liquid Scintillation Analyzer with QuantaSmart TriCarb LCS 3.00 software. ^3^H was measured in the range 2.0-18.6 keV for 5 min on the high sensitivity setting. ^14^C was measure in the range 0.0-156 keV for 2 min on the high-sensitivity setting.

### Ileal ligatures

For ileal ligature surgery in mice, profound anaesthesia was induced by intraperitoneal administration of 100 mg/kg Imalgene1000 (Boehringer-Ingelheim) and 8 mg/kg of Rompun 2% (Bayer). A horizontal laparotomy was performed on the lower abdominal quadrant and a 3-4 cm region of distal ileum retracted and clamped on the extremities using Schwartz vessel clips (World Precision Instruments). Care was taken to avoid compromising the blood supply to the ligated region. Two hundred microlitres of peptidoglycan-AF647 conjugate or PBS control solution was injected into the ligature using a 29G needle. Ileal ligatures were covered gently with gauze soaked with warm DMEM solution (Gibco). After 20 min, the ileal ligature was then collected and the mice sacrificed by cervical dislocation. The excised ileal tissues was opened longitudinally, washed 3 times in PBS and fixed for 3 hours using 4% v/v paraformaldehyde in PBS at 4°C. Tissues were washed 5 times in PBS. Fixed tissues were embedded in 4% LMP agarose (Merck) blocks, then cut into 100 μm-thick sections using a Microm HM 650 V Vibration microtome (Thermo Fisher). Tissue sections were carefully removed from agarose and stored short term in PBS at 4°C in the dark.

### Immunofluorescence imaging

Tissue sections were blocked and permeabilized by incubation in blocking solution, which comprised PBS containing 3% w/v bovine serum albumin (Merck) and 0.4% v/v Triton X-100 (Merck), at 4°C. The following primary antibodies were used: anti-mouse E-cadherin (rat mAb Eccd2, Takara Bio #M108 1:250), anti-mouse Microfold (M) cell (rat mAb NKM 16-2-4, Miltenyi Biotec #130-096-150 1:250), anti-mouse Siglec-F (rat mAb E50-2440, BD Pharmingen #552125, 1:100), Anti-mouse Chromaganin-A (goat pAb sc-1488, Santa Cruz Biotechnology, 1:200). For secondary staining the following antibodies were used: Alexa Fluor 488 goat anti-rat (Invitrogen, 1:500), Alexa-Fluor 488 donkey, anti-goat (Invitrogen, 1:500). For antibody staining, tissues were incubated in blocking solution for 3 hours, then transferred to blocking solution containing primary antibody at the appropriate dilution and incubated overnight at 4°C with gentle agitation. Primary antibody was removed by passage of tissue sections five times in 4 ml of PBS for 5 minutes. Tissue sections were then incubated for 1h at room temperature with secondary antibody and/or other markers as appropriate: Wheat Germ Agglutinin-Alexa Fluor 488 (1:200, Invitrogen), DAPI (1:1000, BD Biosciences), Phalloidin-iFluor 647 (1:200, Abcam). Tissue sections were washed by passage in PBS as before, then mounted on Superfrost Plus microscope slides (Thermo Fisher) using ProLong Gold Antifade reagent (Thermo Fisher) and #1.5 coverslips (VWR). Confocal acquisitions were performed using a Leica TCS SP8 and Leica HyD SP5 confocal microscopes.

### Peptidoglycome purification

Mice were sacrificed and the entire intestinal tract removed. The lumen content of the intestinal tract pressed into 5 mL of PBS and frozen at −80°C. To extract the peptidoglycan, 4 mL of 20% w/v SDS (Interchim) and 11 mL of H_2_O were added directly to the pellet, which was thawed then incubated in a boiling bain-marie for 1 hour with vigorous agitation. After cooling to room temperature, the suspension was filtered through a 30 μm cell strainer, then the standard protocol for purification of peptidoglycan from Gram-positive bacteria was performed, which includes all the steps necessary to purify peptidoglycan from Gram-negative peptidoglycan^40^. The final extract contains purified insoluble peptidoglycan of the gut microbiota. Soluble muropeptides were generated by overnight incubation of the purified extract in 160 μL of 12.5 mM NaH_2_PO_4_ pH 5.6 and 100 U of mutanolysin from *Streptomyces globisporus* ATCC 21553 (Merck) at 37°C. The reaction was stopped by incubation at 100 °C for 10 min, then centrifugated for 5 min at 16,000 xg. The supernatant was collected and stored at −20 °C.

### Mass spectrometry analysis

Muropeptides were reduced by addition of 150 μL of 500 mm borate buffer pH 9, and 50 μL of 20 mg/mL sodium borohydride (NaBH4) solution (Merck) prepared immediately before use. After 30 min incubation at room temperature, the reaction was stopped by adjusting the pH to 4 by addition of 85% orthophosphoric acid (Prolabo). The reaction was centrifugated for 5 min at 16,000g and the supernatant containing reduced muropeptides collected. Ten microlitres of muropeptide solution was diluted 5-fold in mobile phase (formic acid 0.1% in water) before analysis.

Muropeptides analysis was performed by UHPLC-HRMS, using a Dionex Ultimate 3000 UHPLC coupled to a qExactive Focus (Thermo Fisher). Analysis was performed by injecting 10 μL of muropeptides solution onto a Hypersil GOLD C18 aQ C18 (175 Å, 1.9 μm, 2.1 × 150 mm) column, at a temperature of 50 °C at a flow rate of 0.2 mL/min. Mobile phase A: 0.1% formic acid (Optima, Fisher chemical) in water (Optima LC-MS, Fisher chemical); Mobile phase B = 0.1% formic acid in acetonitrile (Optima LC-MS, Fisher chemical). Analytes were separated using a gradient of 0–15% mobile phase B over 30 min. Mass spectrometry was set to positive electrospray ionization mode with a scan range from 200 to 2,500, with full scan data-dependent acquisition of the three most abundant precursor ions for tandem MS by HCD fragmentation. Peaks selection criteria were as follows: peak area threshold = 2,500,000; at least two fragmentation sets; peak intensity >5000.

### HPLC

HPLC analysis of 3H-mesoDAP labelled peptidoglycan was performed using an LC20 Shimadzu HPLC system, equipped with a Hypersil GOLD aQ C18 column (4.6 × 250 mm; Thermo Fisher) at 52 °C, using a flow rate of 0.5 mL/min. Muropeptides were detected at a 206 nm wavelength using a Shimadzu SPD-20A-UV-Vis detector. For collection of muropeptides, peaks were collected from the outlet of the UV detector into 2 mL microfuge tubes. For radioactive measurements of HPLC purified muropeptides, a 10 μL aliquot of muropeptide solution was transferred to a 6 mL scintillation vial and combined with 5 mL of Ultima Gold LLT scintillation cocktail (Perkin Elmer). After equilibration in the dark at room temperature for 4 hours, samples were analysed using a Tri-Carb 3110 TR Liquid Scintillation Analyzer with QuantaSmart TriCarb LCS 3.00 software, with measurement performed in the range 2.0-18.6 keV for 5 min on the high sensitivity setting.

### Data analysis

Scintillation counting data were plotted using Prism GraphPad. Details of statistical analyses performed are indicated on the relevant figure legend.

## References

1. Curciarello, R., Canziani, K. E., Docena, G. H. & Muglia, C. I. Contribution of Non-immune Cells to Activation and Modulation of the Intestinal Inflammation. Front. Immunol. 10, (2019).

2. Chu, H. & Mazmanian, S. K. Innate immune recognition of the microbiota promotes host-microbial symbiosis. Nat Immunol 14, 668–675 (2013).

3. Bouskra, D. et al. Lymphoid tissue genesis induced by commensals through NOD1 regulates intestinal homeostasis. Nature 456, 507–510 (2008).

4. Hergott, C. B. et al. Peptidoglycan from the gut microbiota governs the lifespan of circulating phagocytes at homeostasis. Blood 127, 2460–2471 (2016).

5. Clarke, T. B. et al. Recognition of peptidoglycan from the microbiota by Nod1 enhances systemic innate immunity. Nat Med 16, 228–231 (2010).

6. Arentsen, T. et al. The bacterial peptidoglycan-sensing molecule Pglyrp2 modulates brain development and behavior. Mol Psychiatry 22, 257–266 (2017).

7. Schrijver, I. A. et al. Reduced systemic IgG levels against peptidoglycan in rheumatoid arthritis (RA) patients. Clin Exp Immunol 123, 140–146 (2001).

8. Huang, Z. et al. Antibody neutralization of microbiota-derived circulating peptidoglycan dampens inflammation and ameliorates autoimmunity. Nat Microbiol 4, 766–773 (2019).

9. Molinaro, R., Mukherjee, T., Flick, R., Philpott, D. J. & Girardin, S. E. Trace levels of peptidoglycan in serum underlie the NOD-dependent cytokine response to endoplasmic reticulum stress. J Biol Chem 294, 9007–9015 (2019).

10. Gabanyi, I. et al. Bacterial sensing via neuronal Nod2 regulates appetite and body temperature. Science 376, eabj3986.

11. Bastos, P. A. D., Wheeler, R. & Boneca, I. G. Uptake, recognition and responses to peptidoglycan in the mammalian host. FEMS Microbiol. Rev. (2020) doi:10.1093/femsre/fuaa044.

12. Smith, D. E., Clémençon, B. & Hediger, M. A. Proton-coupled oligopeptide transporter family SLC15: physiological, pharmacological and pathological implications. Mol Aspects Med 34, 323–336 (2013).

13. Wang, Z.-M. et al. Human peptidoglycan recognition protein-L is an N-acetylmuramoyl-L-alanine amidase. J. Biol. Chem. 278, 49044–49052 (2003).

14. Tomasić, J., Ladesić, B., Valinger, Z. & Hrsak, I. The metabolic fate of 14C-labeled peptidoglycan monomer in mice. I. Identification of the monomer and the corresponding pentapeptide in urine. Biochim. Biophys. Acta 629, 77–82 (1980).

15. Kulkarni, D. H. et al. Goblet cell associated antigen passages support the induction and maintenance of oral tolerance. Mucosal Immunology 13, 271–282 (2020).

16. Knoop, K. A., McDonald, K. G., McCrate, S., McDole, J. R. & Newberry, R. D. Microbial sensing by goblet cells controls immune surveillance of luminal antigens in the colon. Mucosal Immunol 8, 198–210 (2015).

17. McDole, J. R. et al. Goblet cells deliver luminal antigen to CD103+ dendritic cells in the small intestine. Nature 483, 345–349 (2012).

18. Miller, M. J., Knoop, K. A. & Newberry, R. D. Mind the GAPs: insights into intestinal epithelial barrier maintenance and luminal antigen delivery. Mucosal Immunol 7, 452–454 (2014).

19. Muise, E. D., Gandotra, N., Tackett, J. J., Bamdad, M. C. & Cowles, R. A. Distribution of muscarinic acetylcholine receptor subtypes in the murine small intestine. Life Sciences 169, 6–10 (2017).

20. Bu, H.-F., Wang, X., Tang, Y., Koti, V. & Tan, X.-D. Toll-like receptor 2-mediated peptidoglycan uptake by immature intestinal epithelial cells from apical side and exosome-associated transcellular transcytosis. Journal of Cellular Physiology 222, 658–668 (2010).

21. Wang, X. et al. Cardiomyocytes mediate anti-angiogenesis in type 2 diabetic rats through the exosomal transfer of miR-320 into endothelial cells. Journal of Molecular and Cellular Cardiology 74, 139–150 (2014).

22. Brugiroux, S. et al. Genome-guided design of a defined mouse microbiota that confers colonization resistance against Salmonella enterica serovar Typhimurium. Nat Microbiol 2, 1–12 (2016).

23. Ladesić, B., Tomasić, J., Kveder, S. & Hrsak, I. The metabolic fate of 14C-labeled immunoadjuvant peptidoglycan monomer. II. In vitro studies. Biochim Biophys Acta 678, 12–17 (1981).

24. Valinger, Z., Ladesić, B., Hrsak, I. & Tomasić, J. Relationship of metabolism and immunostimulating activity of peptidoglycan monomer in mice after three different routes of administration. Int J Immunopharmacol 9, 325–332 (1987).

25. Wolf, A. J. et al. Hexokinase Is an Innate Immune Receptor for the Detection of Bacterial Peptidoglycan. Cell 166, 624–636 (2016).

26. Read, C. B. et al. Cutting Edge: identification of neutrophil PGLYRP1 as a ligand for TREM-1. Journal of Immunology (Baltimore, Md.: 1950) 194, 1417–1421 (2015).

27. Bersch, K. L. et al. Bacterial Peptidoglycan Fragments Differentially Regulate Innate Immune Signaling. ACS Cent. Sci. 7, 688–696 (2021).

28. Zhang, Q. et al. Intestinal lysozyme liberates Nod1 ligands from microbes to direct insulin trafficking in pancreatic beta cells. Cell Res 29, 516–532 (2019).

29. Laman, J. D., Hart, B. A. ’t, Power, C. & Dziarski, R. Bacterial Peptidoglycan as a Driver of Chronic Brain Inflammation. Trends in Molecular Medicine 26, 670–682 (2020).

30. Tosoni, G., Conti, M. & Diaz Heijtz, R. Bacterial peptidoglycans as novel signaling molecules from microbiota to brain. Current Opinion in Pharmacology 48, 107–113 (2019).

31. Hooper, L. V. et al. Molecular Analysis of Commensal Host-Microbial Relationships in the Intestine. Science 291, 881–884 (2001).

32. Shimada, Y. et al. Commensal Bacteria-Dependent Indole Production Enhances Epithelial Barrier Function in the Colon. PLOS ONE 8, e80604 (2013).

33. Singh, R. et al. Enhancement of the gut barrier integrity by a microbial metabolite through the Nrf2 pathway. Nature Communications 10, 89 (2019).

34. Karczewski, J. et al. Regulation of human epithelial tight junction proteins by Lactobacillus plantarum in vivo and protective effects on the epithelial barrier. American Journal of Physiology. Gastrointestinal and Liver Physiology 298, G851–859 (2010).

35. Hayes, C. L. et al. Commensal microbiota induces colonic barrier structure and functions that contribute to homeostasis. Scientific Reports 8, 14184 (2018).

36. Macho Fernandez, E. et al. Anti-inflammatory capacity of selected lactobacilli in experimental colitis is driven by NOD2-mediated recognition of a specific peptidoglycan-derived muropeptide. Gut 60, 1050–1059 (2011).

37. Powell, J. J. et al. An Endogenous Nanomineral Chaperones Luminal Antigen and Peptidoglycan to Intestinal Immune Cells. Nat Nanotechnol 10, 361–369 (2015).

38. Mengin-Lecreulx, D., Siegel, E. & van Heijenoort, J. Variations in UDP-N-acetylglucosamine and UDP-N-acetylmuramyl-pentapeptide pools in Escherichia coli after inhibition of protein synthesis. Journal of Bacteriology 171, 3282–3287 (1989).

39. Hayashi, K. A rapid determination of sodium dodecyl sulfate with methylene blue. Analytical Biochemistry 67, 503–506 (1975).

40. Wheeler, R., Veyrier, F., Werts, C. & Boneca, I. G. Peptidoglycan and Nod Receptor. in Glycoscience: Biology and Medicine (eds. Taniguchi, N., Endo, T., Hart, G. W., Seeberger, P. H. & Wong, C.-H.) 737–747 (Springer Japan, 2015). doi:10.1007/978-4-431-54841-6_147.

