## Supplementary Figure Legends for "Microbiota-induced active translocation of peptidoglycan across the intestinal barrier dictates its within-host dissemination"

### Contributed equally

$ Contributed equally

* Corresponding authors

#### Supplementary Figure Legends

**Fig. S1. Incorporation of [3H]-mDAP into E. coli PGN.** A) HPLC analysis of typical *E. coli* peptidoglycan preparation used in biodistribution assays. B) CPM [3H]-mDAP incorporation relative to muropeptide abundance. C) Microbiota peptidoglycome analysis by LC-MS. The relative abundance of muropeptides from containing mesoDAP, amidated mesoDAP or lysine at the third amino acid position were identified. The relative abundance of individual muropeptides, averaged for two female littermates, is presented. Color-coded labels correspond to the amino-acid present at position three of the peptide stem. GlcNAc; M, MurNAc; aM, 1,6-anhydroMurNAc; 2-5, peptide stem length; N, Asn; D, Asp.

**Fig. S2. The kinetics of [3H]-PGN biodistribution following *per os* administration in mice.** A) [$^{3}$H]-PGN measured by scintillation counting of dissolved, decolored organs and blood between 2h and 8 h post-gavage. Data are presented as CPM per whole organ, or per tissue fragment (duodenum, ileum, colon). Welch's ANOVA comparing time-point groups, excluding control: Brain $p = 0.0564$; Spleen $p = 0.5902$; Liver $p = 0.0895$; Kidney $p = 0.2958$; Heart $p = 0.5178$; Lung $p = 0.0656$; Fat $p = 0.9250$; Duodenum $p = 0.1740$; Ileum $p = 0.4658$; Colon $p = 0.1766$. Pairwise comparisons to control performed using the Mann–Whitney U test. * $p \leq 0.05$; ** $p \leq 0.005$; *** $p \leq 0.0005$. B) The biodistribution of [$^{3}$H]-GM3, [$^{3}$H]-GM4 and [$^{3}$H]-GM4-GM4, administered *per os* in SPF mice. Data are presented as CPM per whole organ, or per tissue fragment. Pairwise comparison with control performed using unpaired t-test. * $p \leq 0.05$; ** $p \leq 0.0050$; *** $p \leq 0.0005$; $p < 0.0001$. Welch's ANOVA comparing muropeptides groups, excluding control: Brain $p = 0.0564$; Spleen $p = 0.5902$; Liver $p =$

0.0895; Kidney $p$ = 0.2958; Heart $p$ = 0.7861; Lung $p$ = 0.0656; Fat $p$ = 0.0925; Thymus $p$ = 0.8678; Duodenum $p$ = 0.1740; Ileum $p$ = 0.4658; Colon $p$ = 0.1766.

**Fig. S3. Effect of labelling strategy parameters on biodistribution of peptidoglycan.** A) Biodistribution of [$^{3}$H]-mesoDAP amino acid administered to mice *per os*. $^{3}$H]-mesoDAP biodistribution was measured by scintillation counting of dissolved, decolored organs, 2h, 6h and 10 h post-gavage. Data are presented as CPM per whole organ or tissue fragment. Welch's ANOVA comparing time-point groups, excluding control: Brain $p$ = 0.0159; Spleen $p$ = 0.0529; Liver $p$ = 0.2269; Kidney $p$ < 0.0001; Heart $p$ = 0.8868; Lung $p$ = 0.1827; Fat $p$ = 0.0066; Duodenum $p$ = 0.0054; Jejunum $p$ = 0.0056; Ileum $p$ = 0.7670. B) Schematics summary of Radiolabeling methodology, indicating the position of the 3H-labelled mesoDAP and 14C-labelled GlcNAc. C) Biodistribution of [$^{3}$H]-PGN versus [$^{14}$C]-PGN administered to mice *per os*. Mice were gavaged with 400,000 cpm of [$^{3}$H]-PGN or [$^{14}$C]-PGN and scintillation counting performed on dissolved, decolored organs 4h post gavage**.** Data normalized as CPM values per g tissue weight. D) [$^{3}$H]-PGN or [$^{14}$C]-PGN biodistribution data presented as CPM per whole organ, or per tissue fragment without normalization. Pairwise comparison to control performed using the Mann–Whitney U test. * $p \leq 0.05$; ** $p \leq 0.005$; *** $p \leq 0.0005$.

**Fig. S4. Biodistribution of different doses of [$^{3}$H]-PGN following intravenous or intraperitoneal administration, without tissue weight normalization.** A) Mice were administered 40,000 CPM of [3H]-PGN intravenously and biodistribution to organs and tissues measured at 1h, 4h, 8h and 24h post-injection**.** Welch's ANOVA comparing time-point groups, excluding control: Brain $p$ = 0.8823; Spleen $p$ = 0.8644; Liver $p$ = 0.5885; Kidney $p$ < 0.0370; Heart $p$ = 0.9196; Lung $p$ = 0.1599; Fat $p$ = 0.3361; Duodenum $p$ = 0.0758; Ileum $p$ = 0.9186; Colon $p$ = 0.9833. B) Comparison of [3H]-PGN distribution administered intravenously versus gavage. Mice were administered 400,000 CPM of [3H]-PGN intravenously or by gavage, and biodistribution to organs and tissues assessed at 1h (intravenous) or 4h (gavage). C) Biodistribution of [$^{3}$H]-PGN administered intraperitoneally**.** Mice were injected intraperitoneally with 400,000 CPM of [$^{3}$H]-PGN and by scintillation counting performed on the dissolved, decolored organs harvested at 2h and 6h post-gavage. Pairwise comparison with between time-points performed using the Mann–Whitney U test, Brain $p$ = 0.0159; Spleen $p$ = 0.6508; Liver $p$ = 0.5476; Kidney $p$ = 0.0079; Heart $p$ = 0.1667; Lung $p$ = 0.8095; Fat $p$ = 0.0079; Duodenum $p$ = 0.0159; Ileum $p$ = 0.1190; Colon $p$ =

0.9365. Pairwise comparison to control performed using the Mann–Whitney U test. * $p \leq 0.05$; ** $p \leq 0.005$; *** $p \leq 0.0005$.

**Fig. S5. Cellular localization of MDP-rhodamine in the mouse ileal epithelium.** No association with MDP-rho was observed for $CgA^+$ enteroendocrine cells, NKM-16-4-$2^+$ M cells or Siglec-$F^+$ tuft-cells, whereas MDP-rho uptake is observed elsewhere in the same field. Yellow asterisks highlight antibody positive cells in each panel.

**Fig. S6. Regulation of the dissemination of [3H]-PGN across the gut.** A) Muscarinic receptor antagonism suppresses the systemic biodistribution of [3H]-PGN without normalization. SPF mice were administered tropicamide, atropine or vehicle control prior to gavage with $[^3H]$-PGN. Scintillation counting was performed on dissolved, decolored organs. Results are presented as CPM per whole organ, per tissue fragment (duodenum, ileum and colon). B) $[^3H]$-PGN biodistribution from the gut is suppressed by GW4869 treatment. Results are presented as CPM per whole organ, per tissue fragment (duodenum, ileum and colon). Pairwise comparison to vehicle control performed using the Mann–Whitney U test. * $p \leq 0.05$; ** $p \leq 0.005$.

**Fig. S7**. **Dependency of [3H]-PGN biodistribution on microbial colonization status of the host.** A) The biodistribution of $[^3H]$-PGN in germ-free (GF) mice, specific pathogen free (SPF) mice and conventionalized mice (previously GF mice co-housed with SPF for 3 weeks) presented as CPM per whole organ, or per tissue fragment (duodenum, ileum, colon). B) Enumeration of aerobically and anaerobically cultured fecal microbiota from germ-free, conventionalized and SPF mice. Feces were collected immediately prior to gavage with $[^3H]$-PGN. C) The biodistribution of $[^3H]$-PGN in germ-free mice 2h and 8h post gavage. D) Biodistribution of [3H]-PGN in SPF mice treated with a broad-spectrum antibiotic cocktail. E) Biodistribution of $[^3H]$-PGN in SPF mice, OMM12 mice and GF mice presented as CPM per whole organ, or per tissue fragment (duodenum, ileum, colon). A, B, D and E) Pairwise comparisons performed using the Mann–Whitney U test. * $p \leq 0.05$; ** $p \leq 0.005$; *** $p \leq 0.0005$. C) Pairwise comparisons performed using Welch's t-test. * $p \leq 0.05$.
