## Supplementary Figures for "Microbiota-induced active translocation of peptidoglycan across the intestinal barrier dictates its within-host dissemination"

### Supplementary Figure 1

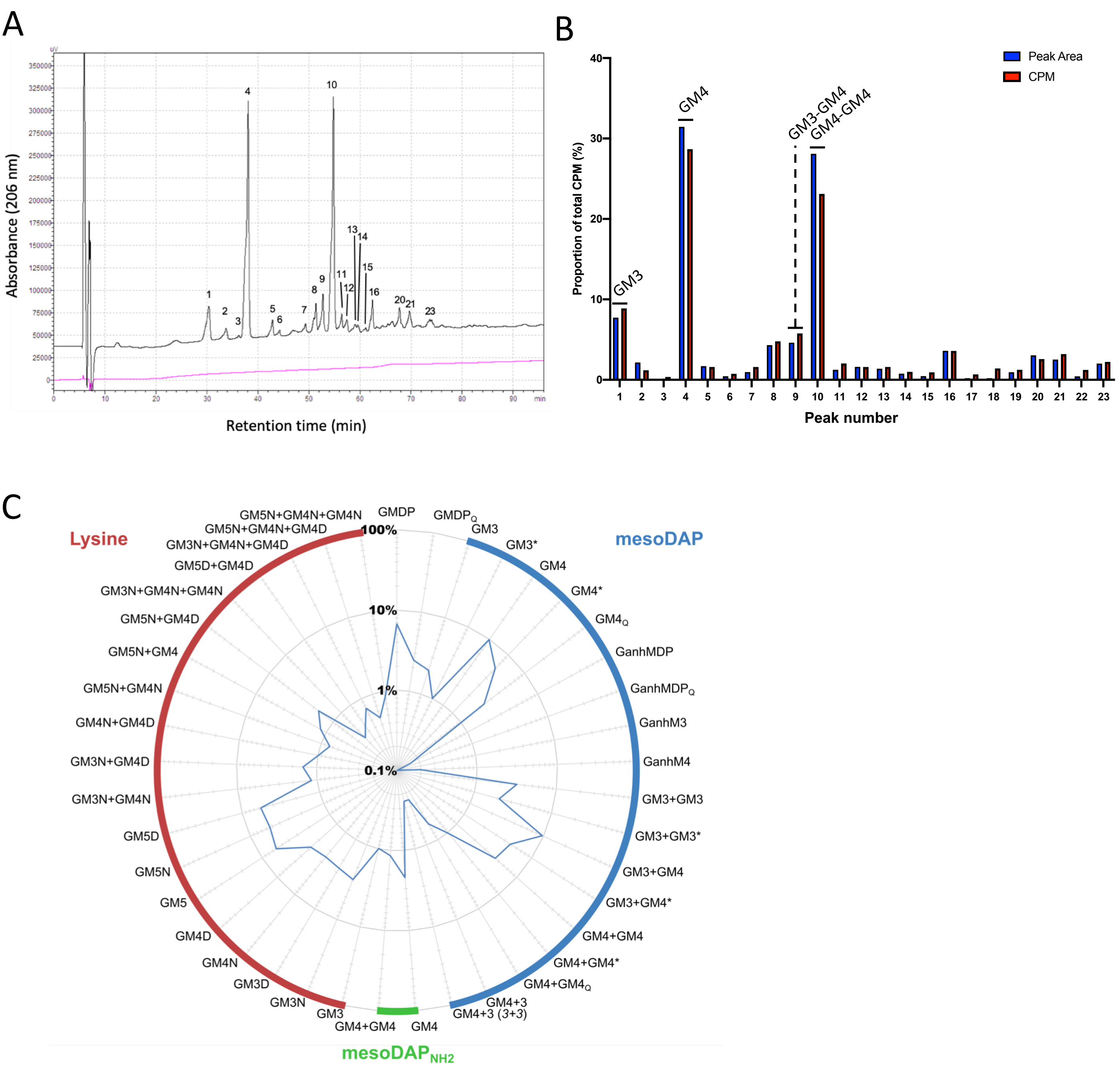

### Supplementary Figure 2

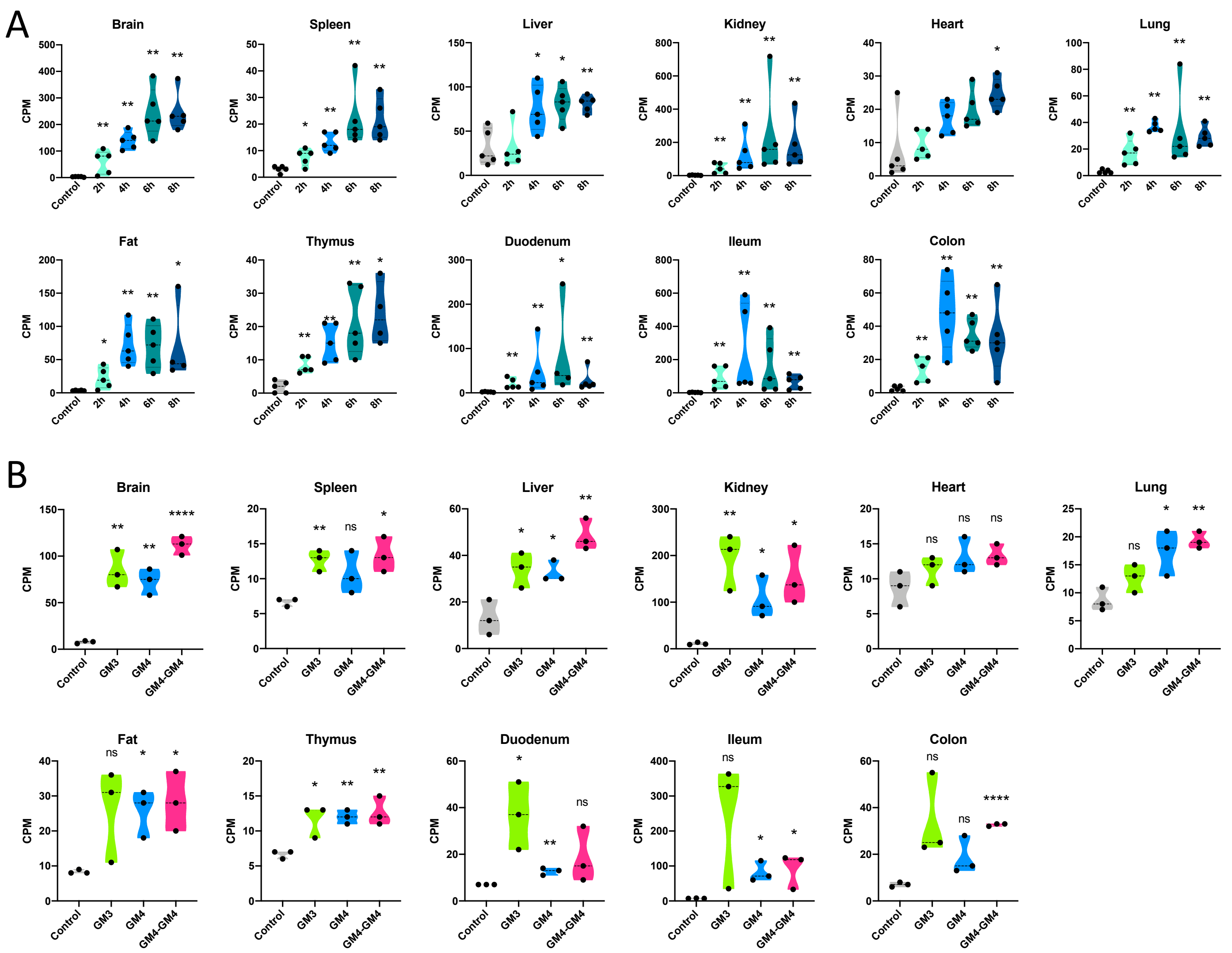

### Supplementary Figure 3

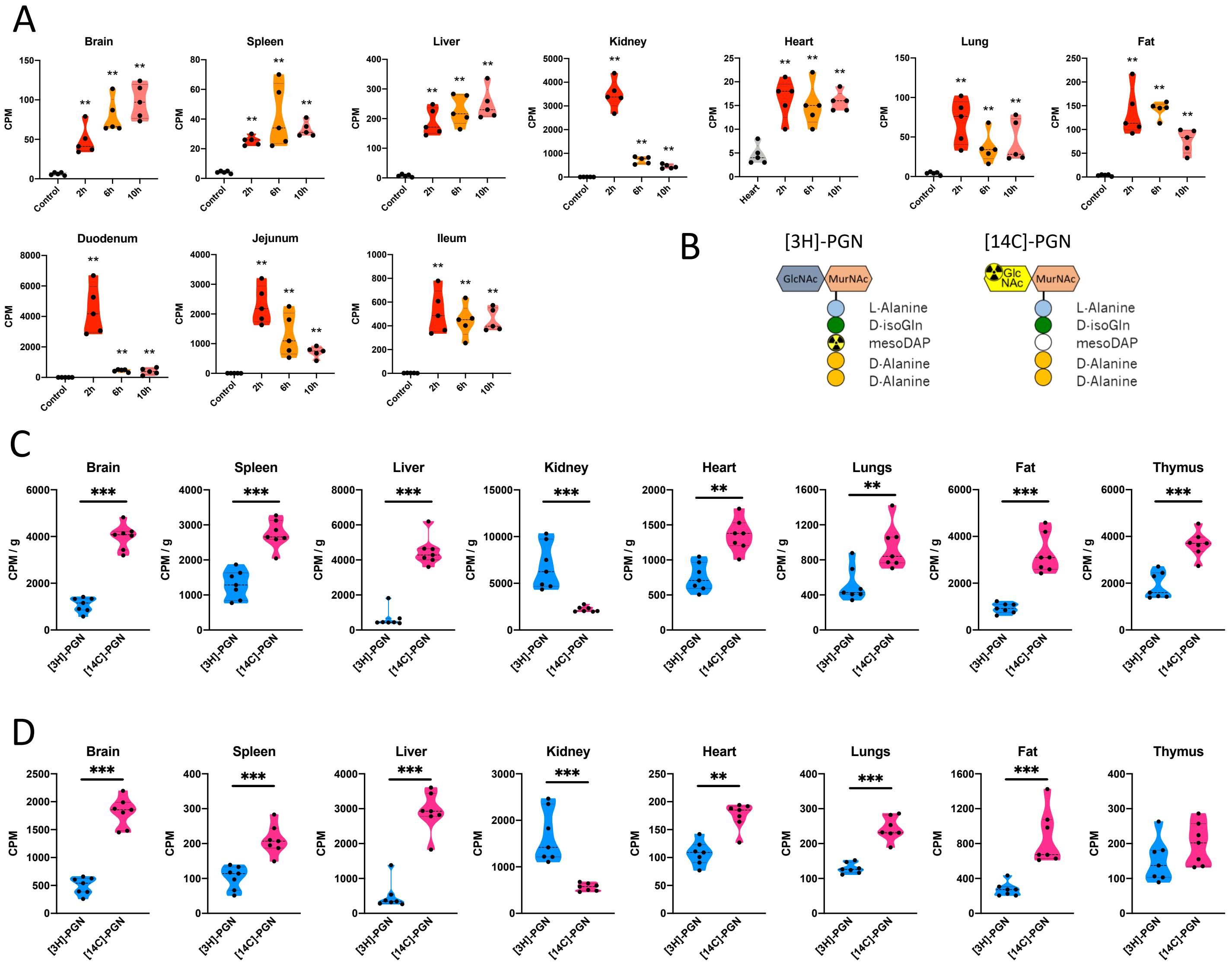

### Supplementary Figure 4

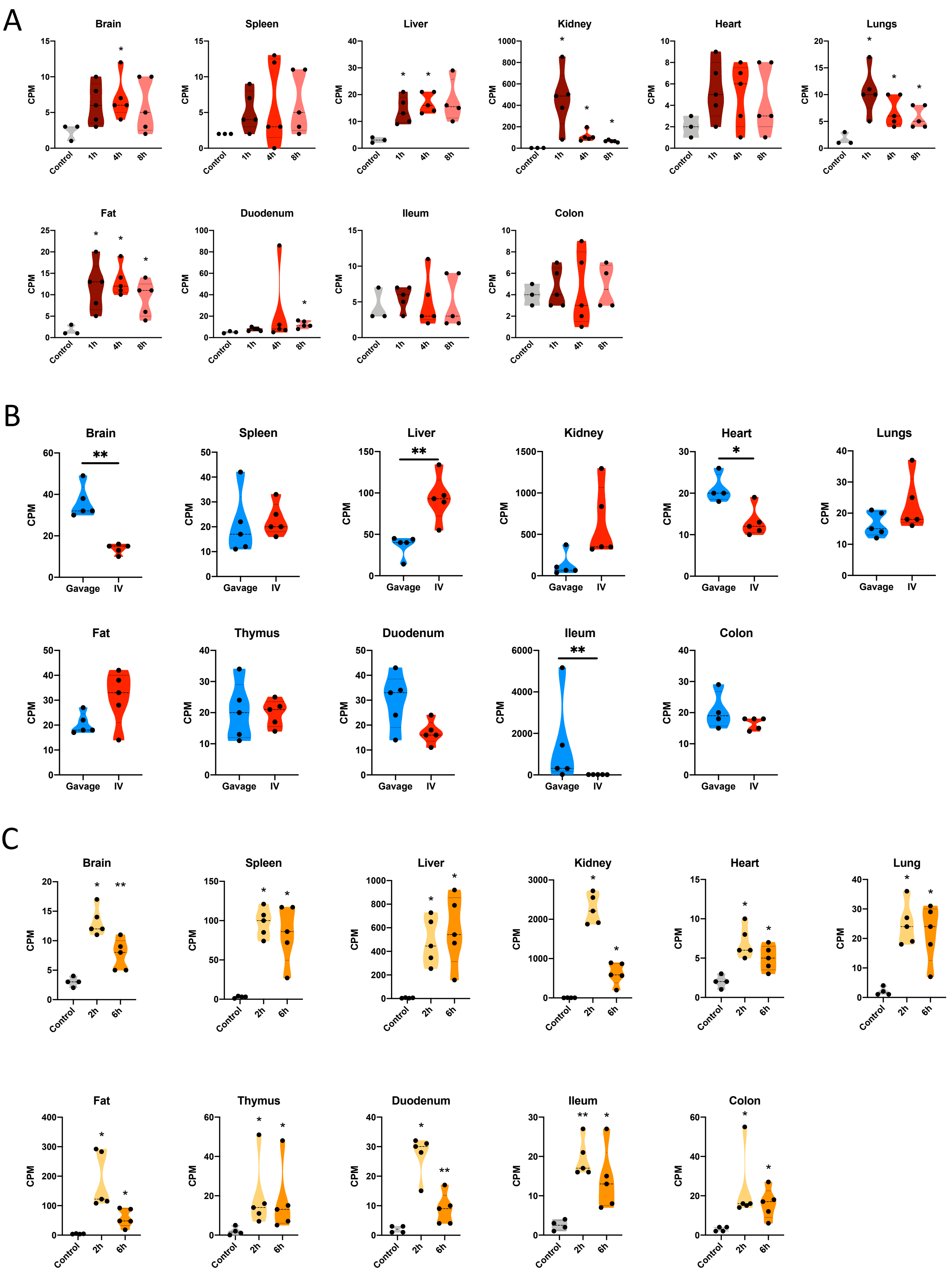

### Supplementary Figure 5

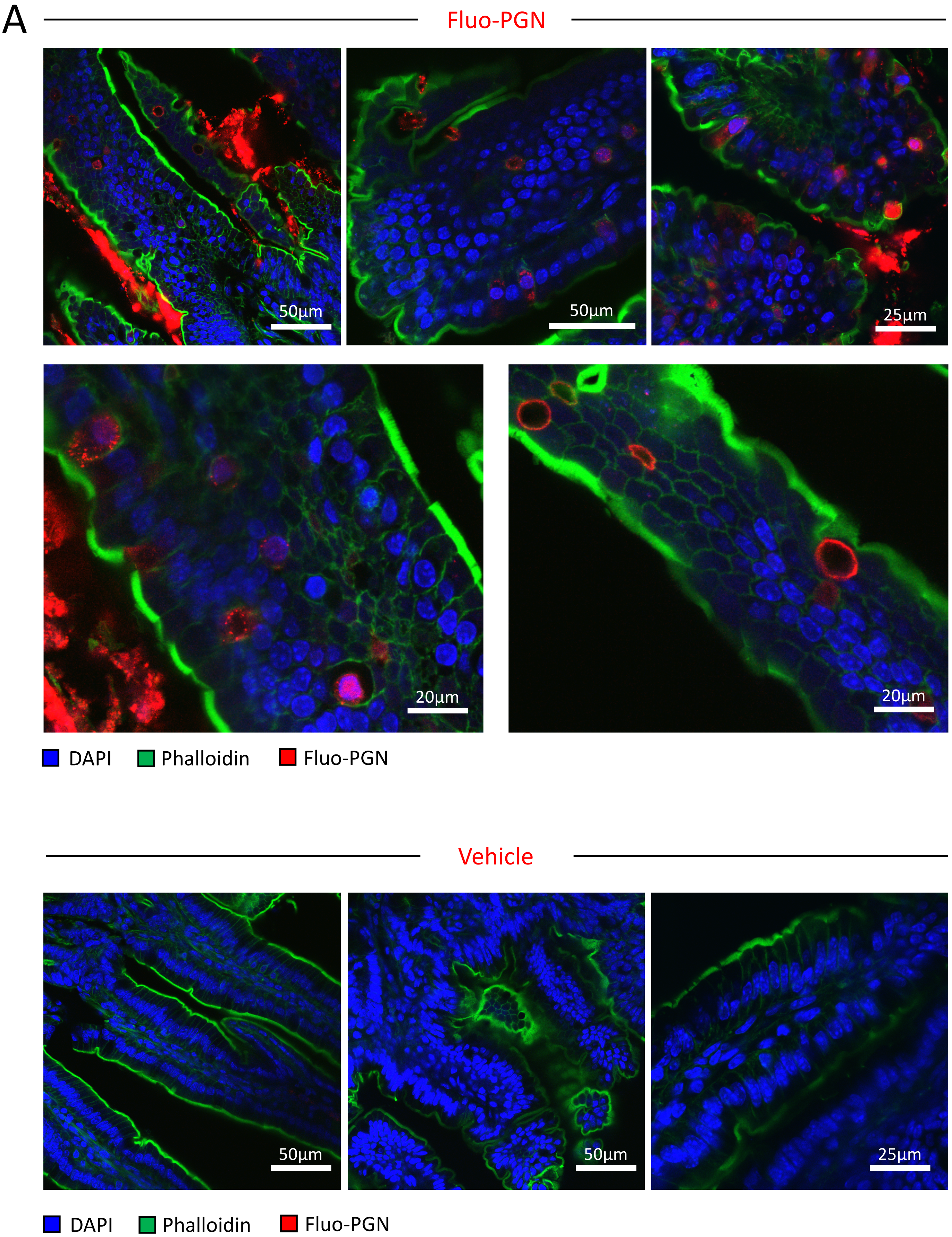

B

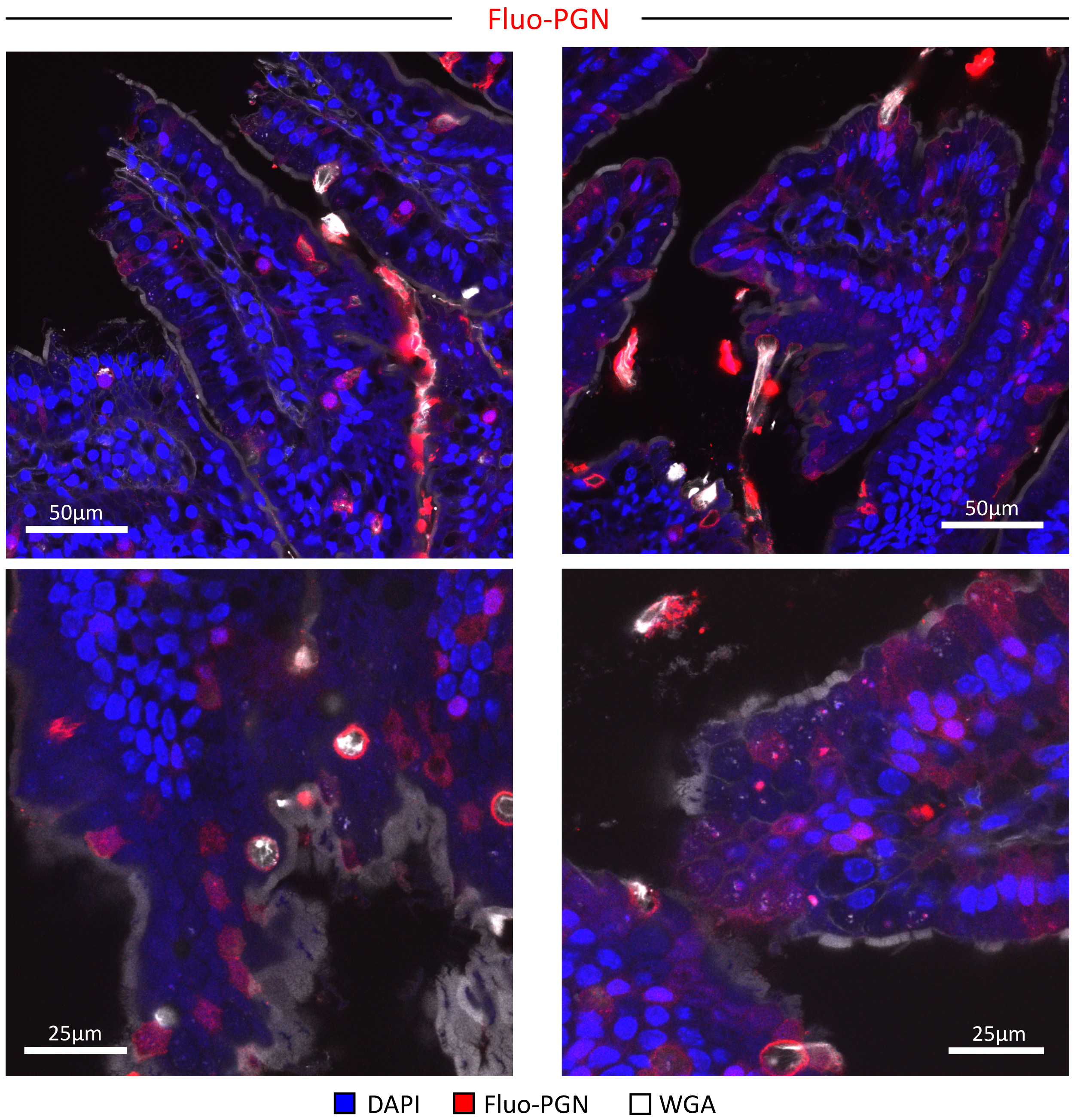
Fluo-PGN
50µm
50µm
25µm
25µm
DAPI
Fluo-PGN
WGA

C

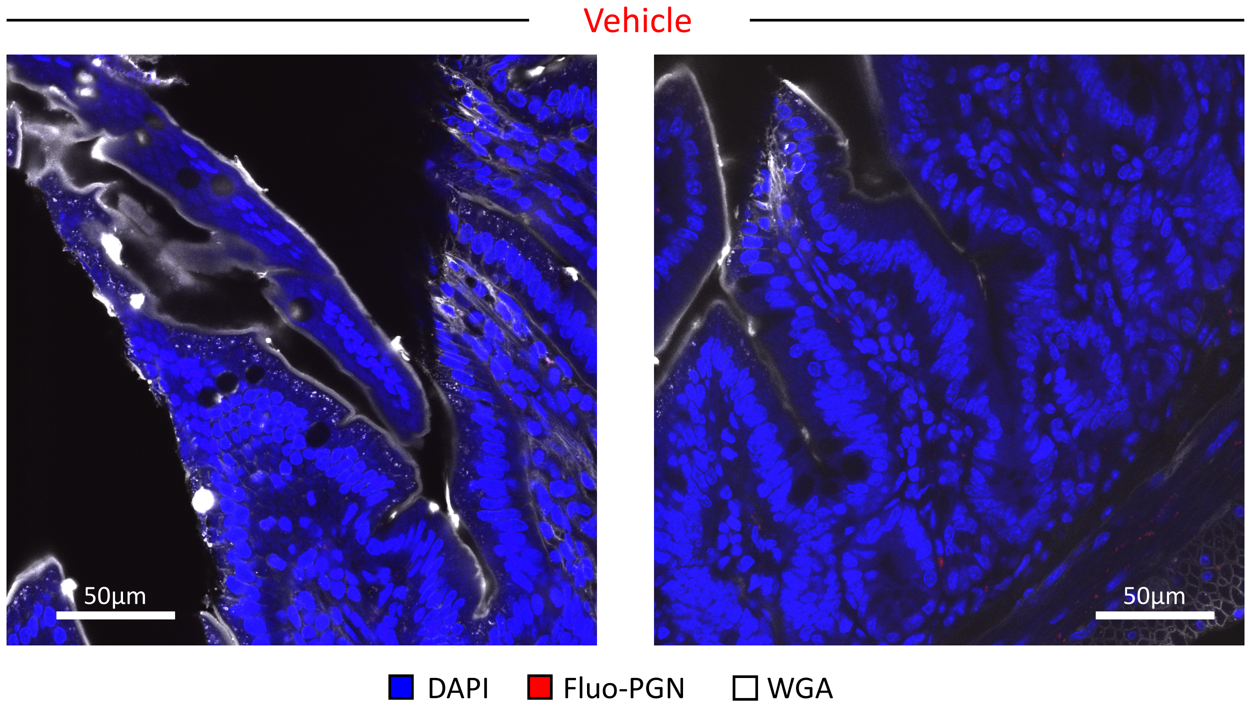
Vehicle
50µm
50µm
DAPI
Fluo-PGN
WGA

D

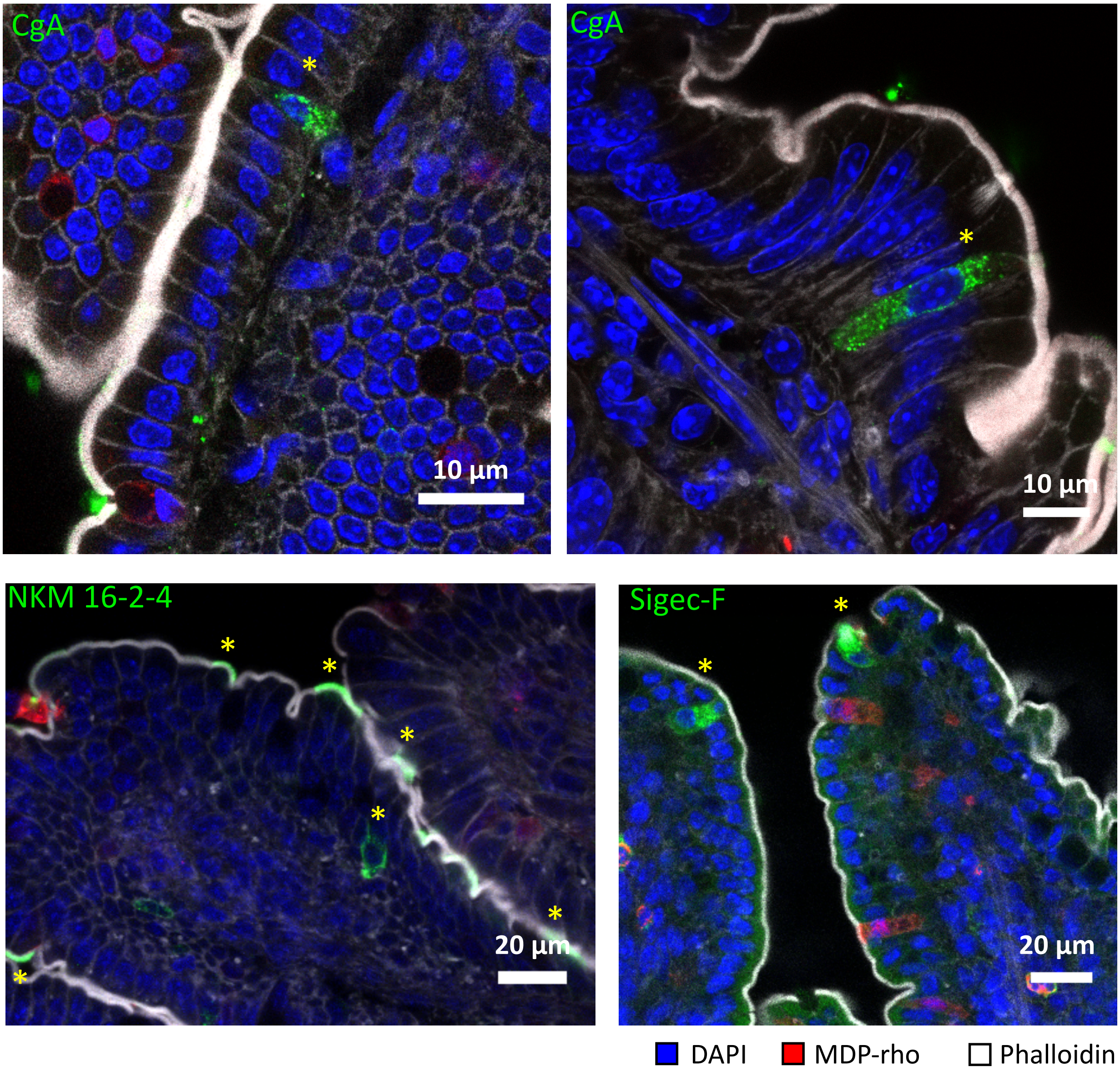
CgA
10 µm
CgA
10 µm
NKM 16-2-4
20 µm
Sigec-F
20 µm
DAPI
MDP-rho
Phalloidin

### Supplementary Figure 6

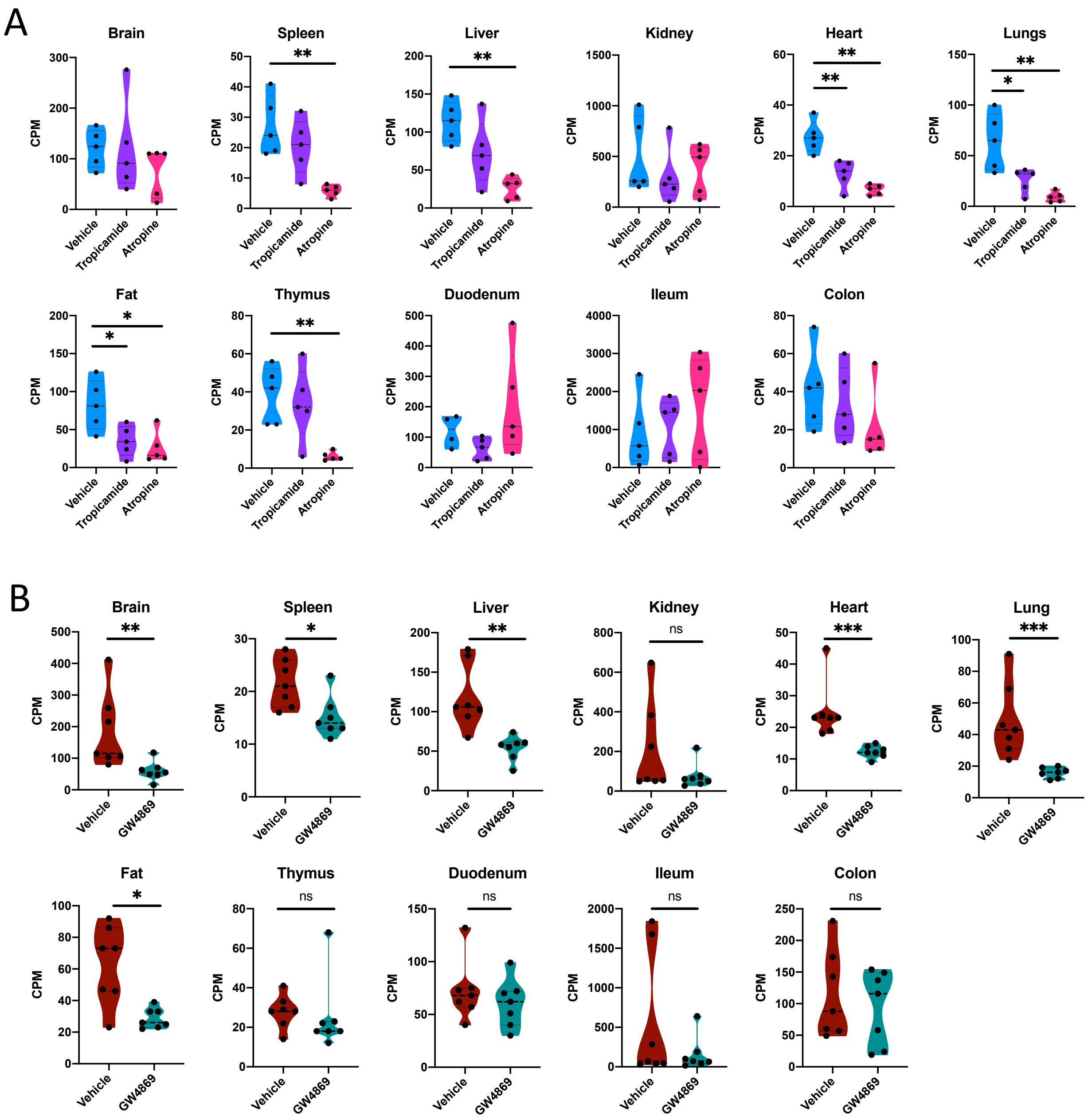

### Supplementary Figure 7

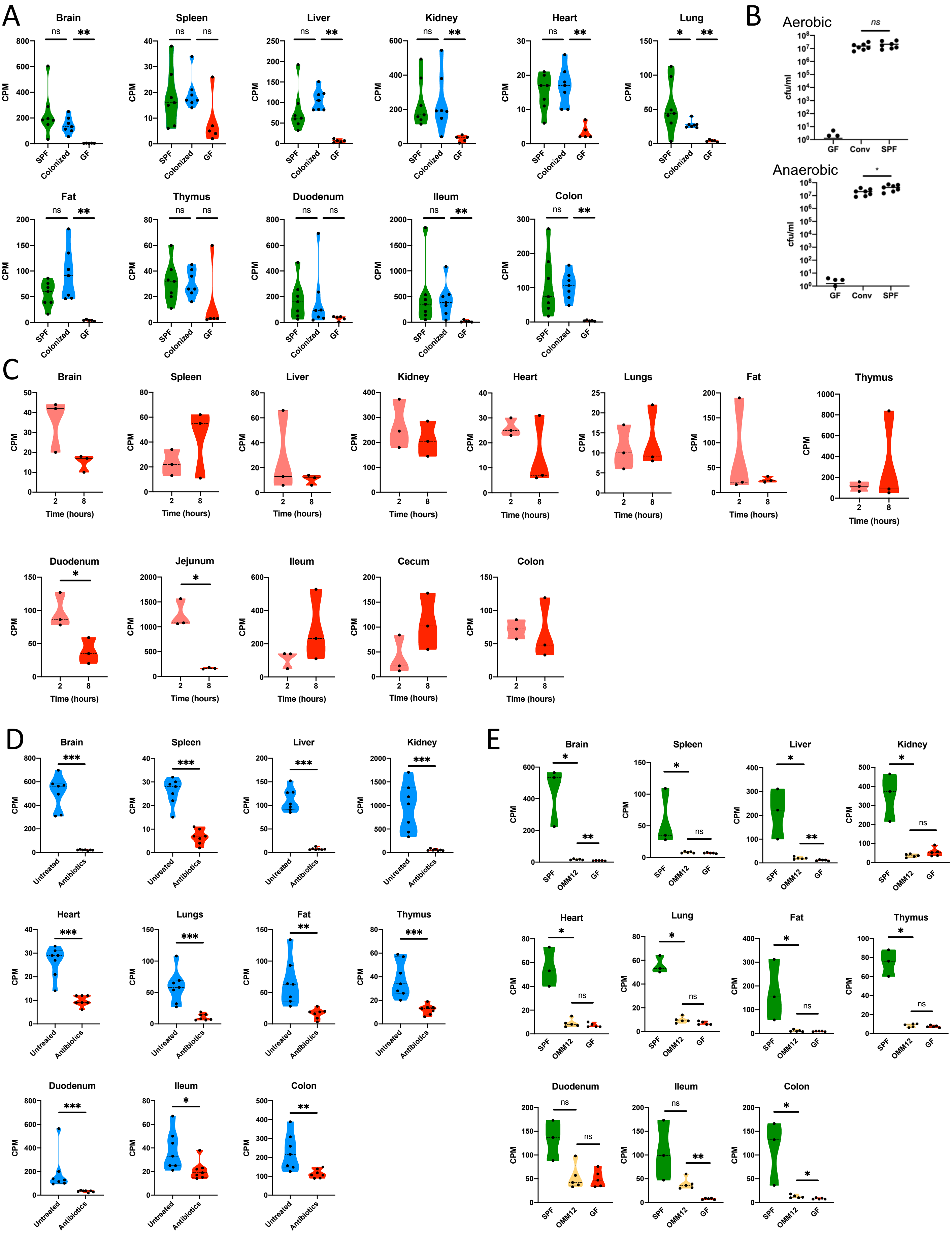
